# Fast computational optimization of TMS coil placement for individualized electric field targeting

**DOI:** 10.1101/2020.05.27.120022

**Authors:** Luis J. Gomez, Moritz Dannhauer, Angel V. Peterchev

**Author notes:** Corresponding Author: Angel V. Peterchev, Address: 40 Duke Medicine Circle, Box 3620 DUMC, Durham, NC 27710, USA.

## Abstract

**Background:** During transcranial magnetic stimulation (TMS) a coil placed on the scalp is used to non-invasively modulate activity of targeted brain networks via a magnetically induced electric field (E-field). Ideally, the E-field induced during TMS is concentrated on a targeted cortical region of interest (ROI).

**Objective:** To improve the accuracy of TMS we have developed a fast computational auxiliary dipole method (ADM) for determining the optimum coil position and orientation. The optimum coil placement maximizes the E-field along a predetermined direction or, alternatively, the overall E-field magnitude in the targeted ROI. Furthermore, ADM can assess E-field uncertainty resulting from precision limitations of TMS coil placement protocols.

**Method:** ADM leverages the electromagnetic reciprocity principle to compute rapidly the TMS induced E-field in the ROI by using the E-field generated by a virtual constant current source residing in the ROI. The framework starts by solving for the conduction currents resulting from this ROI current source. Then, it rapidly determines the average E-field induced in the ROI for each coil position by using the conduction currents and a fast-multipole method. To further speed-up the computations, the coil is approximated using auxiliary dipoles enabling it to represent all coil orientations for a given coil position with less than 600 dipoles.

**Results:** Using ADM, the E-fields generated in an MRI-derived head model when the coil is placed at 5,900 different scalp positions and 360 coil orientations per position (over 2.1 million unique configurations) can be determined in under 15 minutes on a standard laptop computer. This enables rapid extraction of the optimum coil position and orientation as well as the E-field variation resulting from coil positioning uncertainty.

**Conclusion:** ADM enables the rapid determination of coil placement that maximizes E-field delivery to a specific brain target. This method can find the optimum coil placement in under 15 minutes enabling its routine use for TMS. Furthermore, it enables the fast quantification of uncertainty in the induced E-field due to limited precision of TMS coil placement protocols, enabling minimization and statistical analysis of the E-field dose variability.

**Highlights:**

- Auxiliary dipole method (ADM) optimizes TMS coil placement in under 8 minutes
- Optimum coil position is up to 14 mm away from conventional targeting
- Optimum coil orientation is typically near normal to the sulcal wall
- TMS induced E-field is less sensitive to orientation than position errors

## Introduction

Transcranial magnetic stimulation (TMS) is a noninvasive brain stimulation technique [1,2], where a TMS coil placed on the scalp is used to generate a magnetic field that induces an electric field (E-field) in the head. This, in turn, directly modulates the activity of brain regions and network nodes exposed to a high intensity E-field [3,4]. As such, computational E-field dosimetry has been identified by the National Institute of Mental Health as instrumental for determining brain regions stimulated by TMS and for developing rigorous and reproducible TMS paradigms [5]. For efficient and focal stimulation, it is important to position and orient the coil to induce a maximal E-field in the targeted cortical region of interest (ROI) [6,7]. Furthermore, since coil placement protocols have limited precision, it is also important to quantify the variability in the TMS induced ROI Efield due to potential errors in coil placement. This work proposes a novel auxiliary dipole method (ADM) for fast E-field-informed optimal placement of the TMS coil and for quantifying uncertainty in the TMS induced E-field due to possible coil placement errors.

Optimal TMS coil placement is often determined by using scalp landmarks that correlate with targeted cortical ROIs. For example, to stimulate the dorsolateral prefrontal cortex (DLPFC), the coil is often positioned 5 cm anterior to a position that elicits motor evoked potentials in the contralateral hand muscle [8,9]. Alternatively, the coil is centered at a 10–20 coordinate location commonly used for EEG electrode positioning [10,11]. Scalp landmark-based strategies can result in significant misalignments between the coil placement and the targeted cortical ROI [12]. To improve coil placement, MRI imaging data is sometimes used for ‘neuronavigated’ coil positioning. Standard neuronavigated protocols identify the location on the scalp directly over the targeted cortical site’s center of mass as the optimal coil center position [6,7,12]. This is because commonly used TMS coils have a figure-8 winding configuration that generates a primary E-field (E-field in the absence of the subject’s head) that is strongly concentrated underneath its center [13]. However, the optimum coil placement site on the scalp can be shifted up to 12 mm (5.5 mm on average) away from the scalp location directly above the center of mass [14] by the secondary E-field generated inside the subject's head due to charge build-up on tissue interfaces [15–17].

The orientation of the TMS coil is typically chosen so that the direction of the induced E-field is perpendicular to the ROI's sulcal wall. This orientation is known to maximize the magnitude of the E-field in the ROI [14,18,19] and therefore corresponds to the lowest threshold for cortical activation, which is reinforced by the perpendicular orientation of the main axon of pyramidal neurons relative to the sulcal wall [4]. Indeed, E-field directed into the ROI sulcal wall requires, on average, lowest TMS coil currents to evoke a motor potential [20] and is close to the optimal orientation for targeting each region of the hand motor cortical area [14,21–23]. Therefore, in the absence of an explicit model of neural activation, choosing a coil position and orientation that maximizes the E-field strength is a suitable objective.

A further limitation of TMS procedures is that they have a limited, and often unquantified, precision and accuracy of determining and maintaining the coil placement. Even with gold-standard neuronavigation and robotic coil placement, coil position error can exceed 5 mm [24–26]. As such, there is an uncertainty in the TMS induced E-field resulting from uncertainty in the coil placement. This may result in variability in the outcomes of TMS interventions and needs to be quantified. Therefore, in addition to linking the external coil placement and current to the E-field induced in the brain, computational models should ideally account for coil placement uncertainty.

The total E-field induced in cortical ROIs can be determined by using MRI-derived subject-specific volume conductor models and the finite element method (FEM) or the boundary element method (BEM) [23,27–33]. Evaluating the TMS induced E-field for a single coil position using a standard resolution head model currently requires 35 seconds using FEM [34] or 104 seconds using BEM accelerated with the fast-multipole method (FMM) [35]. Presently, E-field-informed optimal coil placement requires iterative execution of such simulations until an optimal coil position and orientation are found out of a large number of possible options [36,37]. As such, computational requirements limit the routine use of E-field-informed optimization of coil placement. For example, for this reason we restricted the individual model-based dosing in a recent TMS study to selecting only the coil current setting [37].

This paper introduces a computational approach that enables fast E-field-informed optimization of coil placement using high resolution individual head models. Our framework is exceptionally computationally efficient as it can evaluate the E-fields generated in a targeted cortical ROI using a standard MRI-derived head model when the coil is placed at 5,900 different scalp positions and 360 coil orientations per position in under 15 minutes using a laptop computer. This approach is based on the observation that to determine the optimum coil placement on the scalp it is unnecessary to evaluate the E-field induced outside the cortical ROI. This enables the use of the electromagnetic reciprocity principle to compute only the E-field induced in the ROI. Reciprocity has been used previously in the context of BEM simulations of TMS induced E-fields [38,39]. These previous uses of reciprocity have two bottlenecks: First, they require the determination of either the magnetic field or E-field due to many electromagnetic point sources, which limits their use to low-resolution head models. Second, they are limited to isotropic head models, and thus they cannot account for brain tissue conductivity anisotropy. Here we avoid the low-resolution bottleneck by using FMM [40], which enables the rapid calculation of fields due to many electromagnetic point sources. Furthermore, to enable the use of anisotropic conductivity head models, we apply reciprocity by using directly conduction currents in the brain [41]. These modifications to previous uses of reciprocity for TMS simulations make our E-field-informed coil placement framework compatible with head models from common transcranial brain stimulation simulation pipelines [17,31]. Finally, we introduce a method for approximating the coil currents by auxiliary dipoles that can represent all coil orientations for a given coil position with less than 600 dipoles. This enables the rapid generation of maps that quantify the dependence of the E-field on both coil position and orientation. Furthermore, we post-process these maps to quantify the Efield uncertainty due to the limited accuracy and precision of coil placement methods.

The rest of this paper is structured as follows. First, we describe the proposed approach to rapidly and optimally position and orient a TMS coil on the scalp to maximize a specific E-field component or the E-field magnitude in a brain ROI. Second, the approach is benchmarked in terms of accuracy relative to analytical solutions of a spherical head model and in terms of runtime and memory requirement relative to results obtained by using FEM directly to determine the TMS induced E-fields. Third, we use a number of detailed realistic head models and target cortical ROIs from TMS experiments to compare the proposed method to a conventional approach that places the coil to minimize its distance to the cortical ROI. Finally, we use this approach to estimate rapidly the E-field variation related to coil placement errors that inevitably occur with any TMS procedure.

Software implementations of the methods developed in this paper are available [42] for use in MATLAB and Python environments enabling easy integration with existing transcranial brain stimulation software packages (e.g., SimNIBS [43,44] and SCIRun [30]) as add-on toolkits.

## Methods

TMS coil placement is specified by the position of its center, **R**, and its orientation, which is defined by a unit vector 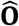 that denotes the direction of the primary E-field directly under the coil center (Fig. 1A). The average E-field along a specified unit vector 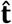 (i.e., 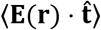, where 〈·* computes the average over the ROI) and the average E-field magnitude (i.e., 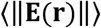 in the ROI) are both functions of coil position and orientation. We formulate the goal of optimal TMS coil placement to be the determination of a coil position **R_opt_** and orientation 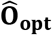 that maximizes either 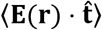 or 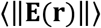. For example, 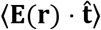 can be maximized if a preferential field direction 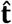 for activating the targeted neural population is known, whereas 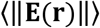 is maximized whenever this information is missing. For clarity, we sometimes emphasize the dependence of 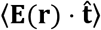 on coil position and orientation by referring to 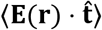 as 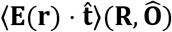.

**Figure 1.**
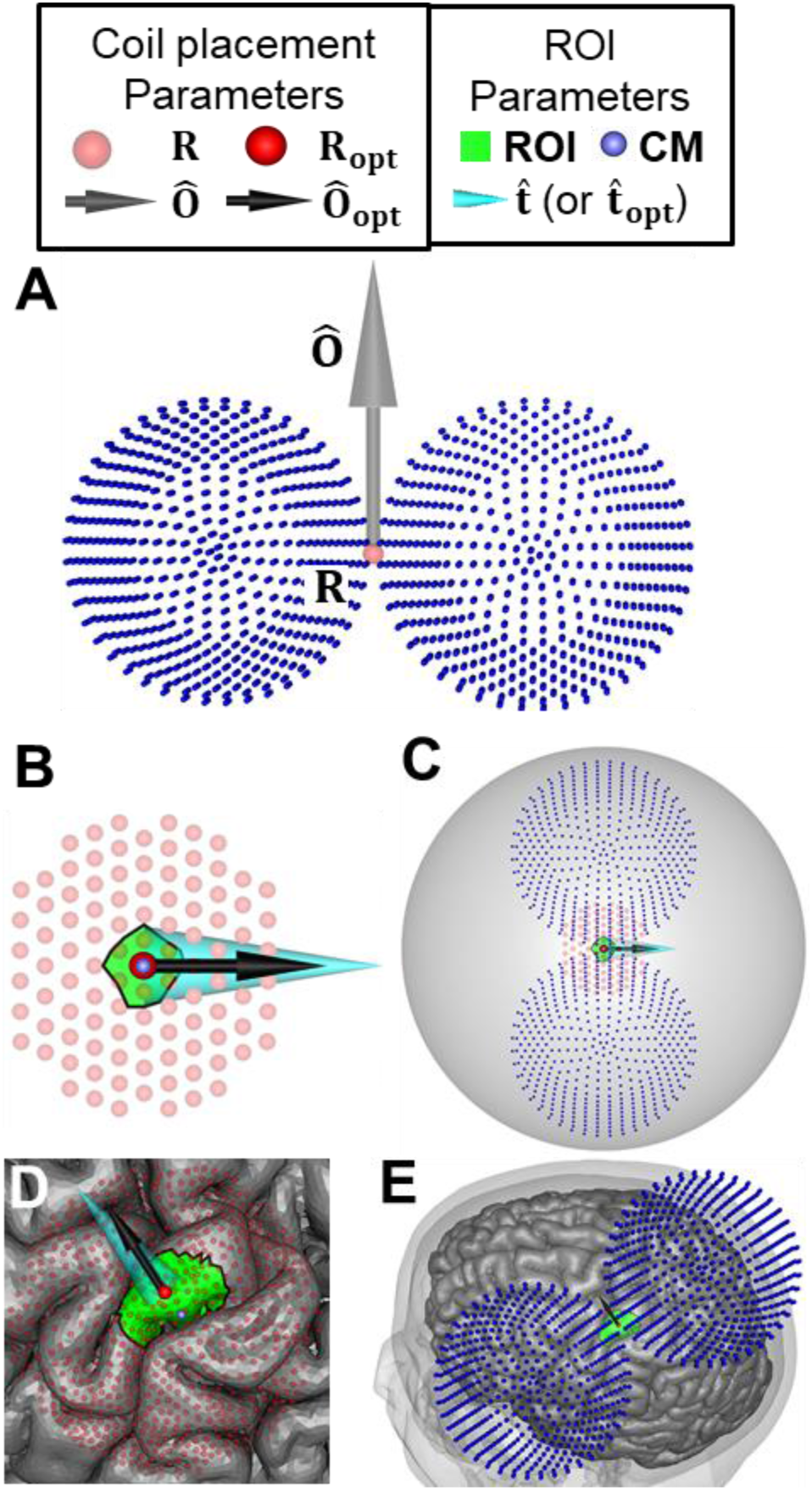
Conventions for describing TMS coil placement in this paper. (A) Figure-8 coil placement is defined by position **R** and orientation 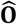 (the dipoles representing the coil model are shown as small dark blue dots). (B) Pink dots indicate candidate coil positions above the brain ROI colored in green. The ROI center of mass (CM) is indicated by a small light blue dot. The optimal coil position and orientation are indicated by a red dot and black arrow, respectively. Wide cyan arrow indicates either the preferred E-field direction 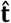 in the ROI, if it is specified, or the average E-field direction 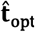 in the ROI resulting from maximization of the E-field magnitude. (C) For the spherical head model, the optimum coil position is directly above the ROI CM and its orientation is aligned exactly with the specified 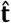. (D,E) The same concepts illustrated for an MRI-based head model (Ernie). For such models, the optimum coil position and orientation can differ from CM and 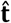 (or 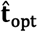), respectively.

We describe a fast reciprocity-based method for evaluating 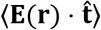. This method is used for rapid determination of **R_opt_** and 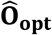 from a large number of candidate coil positions and orientations. Specifically, we define **R_opt_** and 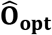 as the candidate position and orientation that maximize 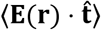 (Figs. 1B-E). Furthermore, this method is extended to find **R_opt_** and 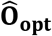 that maximize 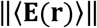. For typical ROIs, 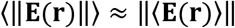. Therefore, the maximization of 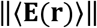 also results in the maximization of 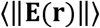. This enables ADM to maximize the E-field magnitude in an ROI. The average E-field direction in the ROI associated with the optimal coil placement, 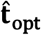, emerges from this optimization as well.

### Reciprocity method for determining the average E-field along a predetermined direction

Electromagnetic reciprocity is an equivalence relationship between two scenarios (Fig. 2). In one scenario, the TMS coil, modeled as impressed electric current **J**(**r**; *t*) = *p*(*t*)**J**(**r**), generates an Efield **E**(**r**; *t*) = *p*’(*t*)**E**(**r**) inside the head (Fig. 2A). Here *t* is time, **r** is a Cartesian location, and *p*(*t*) and *p*’(*t*) are the temporal variation of the TMS coil current and its derivative, respectively. In the second scenario, a source **J_c_**(**r**; *t*) = *p*(*t*)**J_c_**(**r**) inside the head induces an E-field **E_c_**(**r**; *t*) = *p*’(*t*)**E_c_**(**r**) in the TMS coil (Fig. 2B). Reciprocity dictates that the reaction integral between **E**(**r**; *t*) and **J_c_**(**r**; *t*) is equal to the reaction integral between **J**(**r**; *t*) and **E_c_**(**r**; *t*),

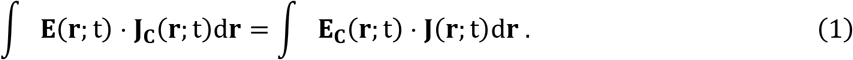

**Figure 2.**
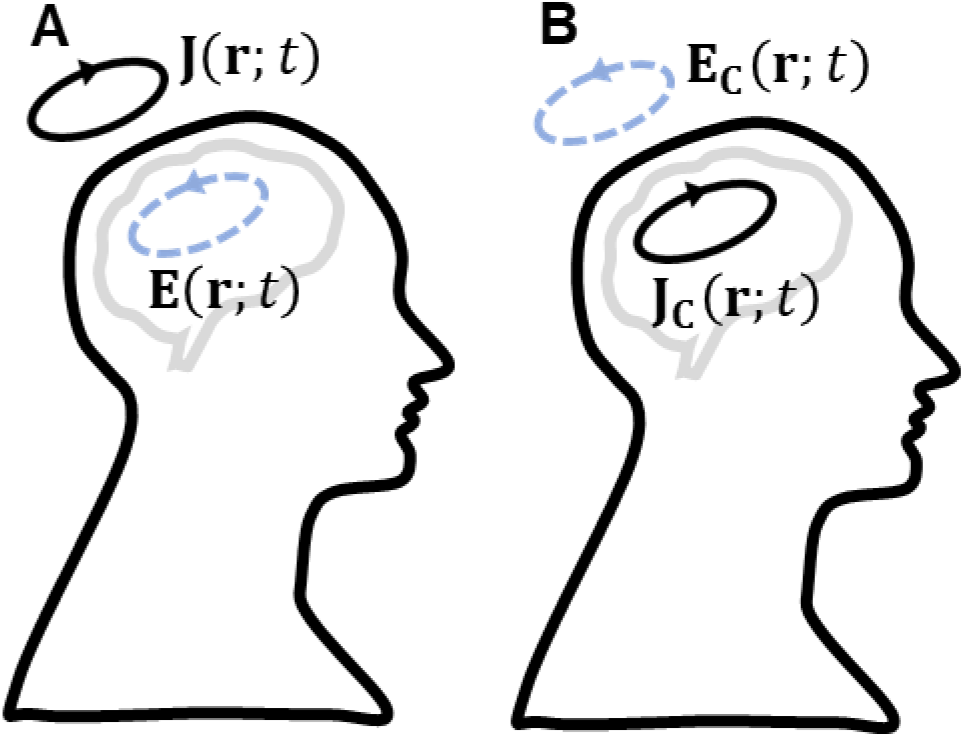
Reciprocal scenarios: (A) The TMS coil current generates an E-field inside the brain. (B) A brain current source generates an E-field where the coil resides.

Here we choose **J_c_**(**r**; *t*) = 0 outside of the ROI and 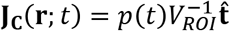 inside it, where *V_ROI_* is the volume of the ROI. Reciprocity results in the average E-field along 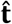 in the ROI being equal to

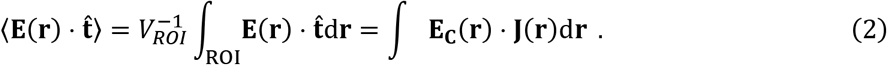

Since for TMS pulse waveform frequencies the E-field spatial and temporal components are separable (quasi-stationary) [45], the temporal variation of all currents and E-fields— *p*(*t*) and *p*’(*t*), respectively—is omitted in this and subsequent notation.

Computing 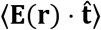 using Eq. (2) requires the evaluation of **E_c_**(**r**) outside of the conductive head. This is done by computing the primary E-field due to the cortical current **J_c_**(**r**) as well as the secondary E-field due to the conduction currents 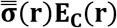, where 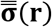 is the conductivity tensor inside the head. Thus

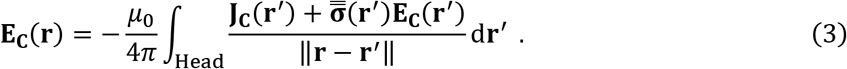

Here *μ*_0_ is the permeability of free space, and the integration is done over the head domain. In the following section, we describe the FEM approach we used to compute **E_c_**(**r**) inside the head, and numerical integration rules used to estimate Eqs. (2)–(3).

### Discretization of the reaction integral

The E-field inside the head due to **J_c_**(**r**) satisfies Laplace’s equation, 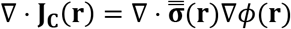, where **E_c_**(*r*) = ∇*ϕ*(**r**). To solve for *ϕ* and ∇*ϕ* we used an in-house 1^nd^ order FEM method [33]. First, the head model is discretized into a tetrahedral mesh having *P* nodes and *N* tetrahedrons, and each tetrahedron is assigned a constant tissue conductivity tensor. Second, the scalar potential *ϕ* is approximated by piece-wise linear nodal elements *N_m_*(*r*) (where *m* = 1,2,…,*P*) [46]. Third, weak forms of Laplace’s equation are sampled also using piecewise-linear nodal elements as testing functions in a standard Galerkin procedure. This results in

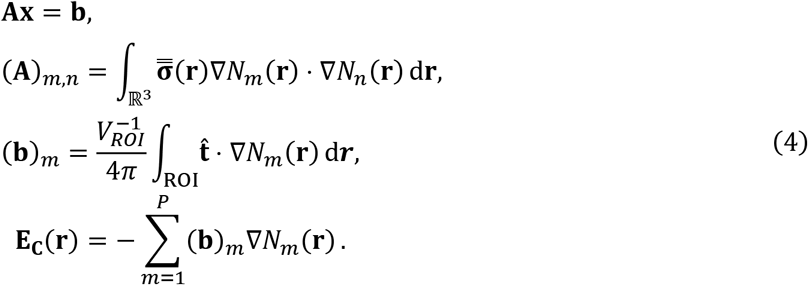

Entries (**A**)_*m,n*_ are computed analytically using expressions provided in [46] and ROI denotes the support of **J_c_**(**r**). The system of equations in Eq. (4) is solved using a transpose-free quasi-minimal residual iterative solver [47] to a relative residual of 10^−10^.

Samples of **E_c_**(**r**) outside the head are obtained via Eq. (3) and using the FEM solution. The volume cortical and conduction currents are approximated by current dipoles on the centroid of each tetrahedron. The current dipoles are computed as 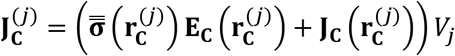, where 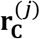 is the centroid, and *V_j_* is the volume of the *j*^th^ tetrahedron. When the above conduction current approximation is applied to Eq. (3), it yields

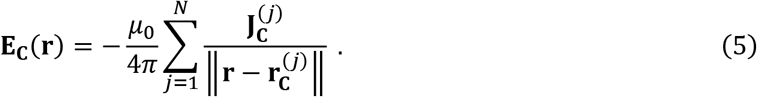

Typically coil models consist of dipoles 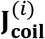 at locations 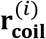, where *i* = 1,2, …,*M*. As a result, using a coil dipole model and Eq. (5) to evaluate Eq. (2) results in

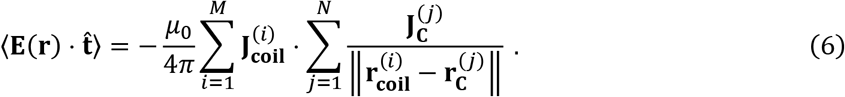

Evaluating Eq. (5) at a single position requires evaluating the sum of *N* entries (i.e., the number of computations scales with the number of tetrahedrons). Furthermore, computing 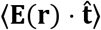 using Eq. (6) requires the evaluation of Eq. (5) at *M* positions (i.e., the number of computations scales with the number of tetrahedrons times the number of coil model dipoles *M × N*). If one wants to evaluate 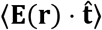 for *L* different coil placements, the number of computations will scale as *L×M×N*. As a result, there are trade-offs between how large *L, M*, and *N* can be while maintaining a tractable number of computations. For example, if the number of tetrahedrons is large we are limited in the number of coil placements for which we can evaluate the E-field. Conversely, if we would like to evaluate the E-field for many coil placements, the resolution of the head model has to be lowered. This is a common limitation of reciprocity-based E-field BEM solvers for TMS [38,39]. To lower the computational cost, we use FMM, which was designed to lower the computational complexity of the calculation of Eq. (6). Specifically, it reduces the total computation from scaling as *N×M×L* to scaling as the maximum between *N* and *M×L*. Using FMM enables us to compute Eq. (6) with both dense head meshes (i.e., large *N*) and for many coil placements (i.e., large *M × L*). Moreover, we generate an auxiliary dipole model, described in the following sections, that leverages the same set of dipole locations for multiple coil orientations to speed up our method further. Note that the method as presented here does not work with magnetic dipole TMS coil models. Its extension to magnetic sources requires minor modifications given in the supplemental material. We created software implementations for both electric and magnetic dipole sources, available online [42].

### Optimizing E-field magnitude

Oftentimes, it is desired to maximize the average E-field magnitude in the ROI

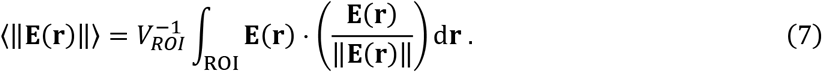

The E-field direction changes slowly as a function of position, as such, it can be well approximated as unidirectional. For a given coil position **R** and 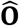 the best possible approximation of 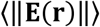 that can be obtained while replacing unit vector 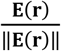 by a constant unit-vector (i.e. unidirectional approximation) is 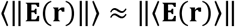. In other words, instead of computing the magnitude and then taking the average over the ROI, the component-wise average of the vector is first computed and then the magnitude is taken. The unidirectional approximation will be less accurate for larger ROIs relative to smaller ones. To assess the accuracy of the unidirectional approximation as a function of ROI size, we compared results obtained by directly computing 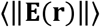 with ones for 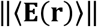 for ROIs of varying sizes. Furthermore, we ran SimNIBS [48] coil placement optimization to maximize 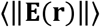 and compared with our results obtained by maximizing 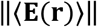.

To use ADM to compute 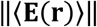. For each candidate coil position and orientation, the ADM is run three times to compute 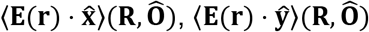, and 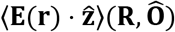. From these three principal directions we compute 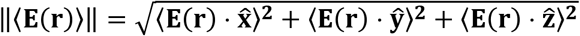. Note that having the three principal directions also enables us to compute 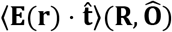 for any 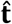 rapidly using the linear decomposition 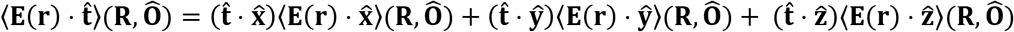. The linear decomposition indicates that computing 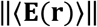 is equal to computing 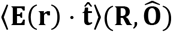 while choosing 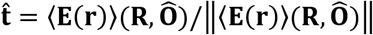. As such, maximizing 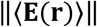 is equal to maximizing 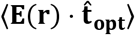, where 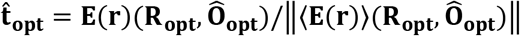. In other words, the E-field magnitude optimization also produces a vector that indicates the direction along which E-field has to be maximized to also maximize E-field magnitude.

### Auxiliary dipole approximation

Here we describe the procedure used to approximate all coil orientations using a few dipoles located at auxiliary nodes (Fig. 3). The dipoles are chosen as Gauss-Legendre tensor product nodes of a rectangular box enclosing the coil (Fig. 3B). Assume that the x-y axis is chosen as the coil plane, and it consists of *M* dipoles 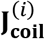 at locations 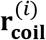, where i = 1,2, …*M* (Fig. 3A). From 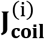 and 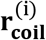, we generate *P* coil dipole models 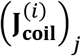 at locations 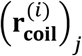, where *j* = 1,2, …*P*, by rotating 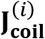 and 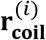 (Fig. 3B). Assume that all the coil models can be bound by a box 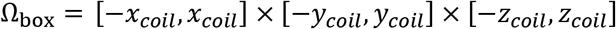 outside the head. For each coil orientation, we need to compute

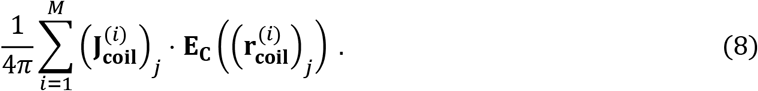

**Figure 3.**
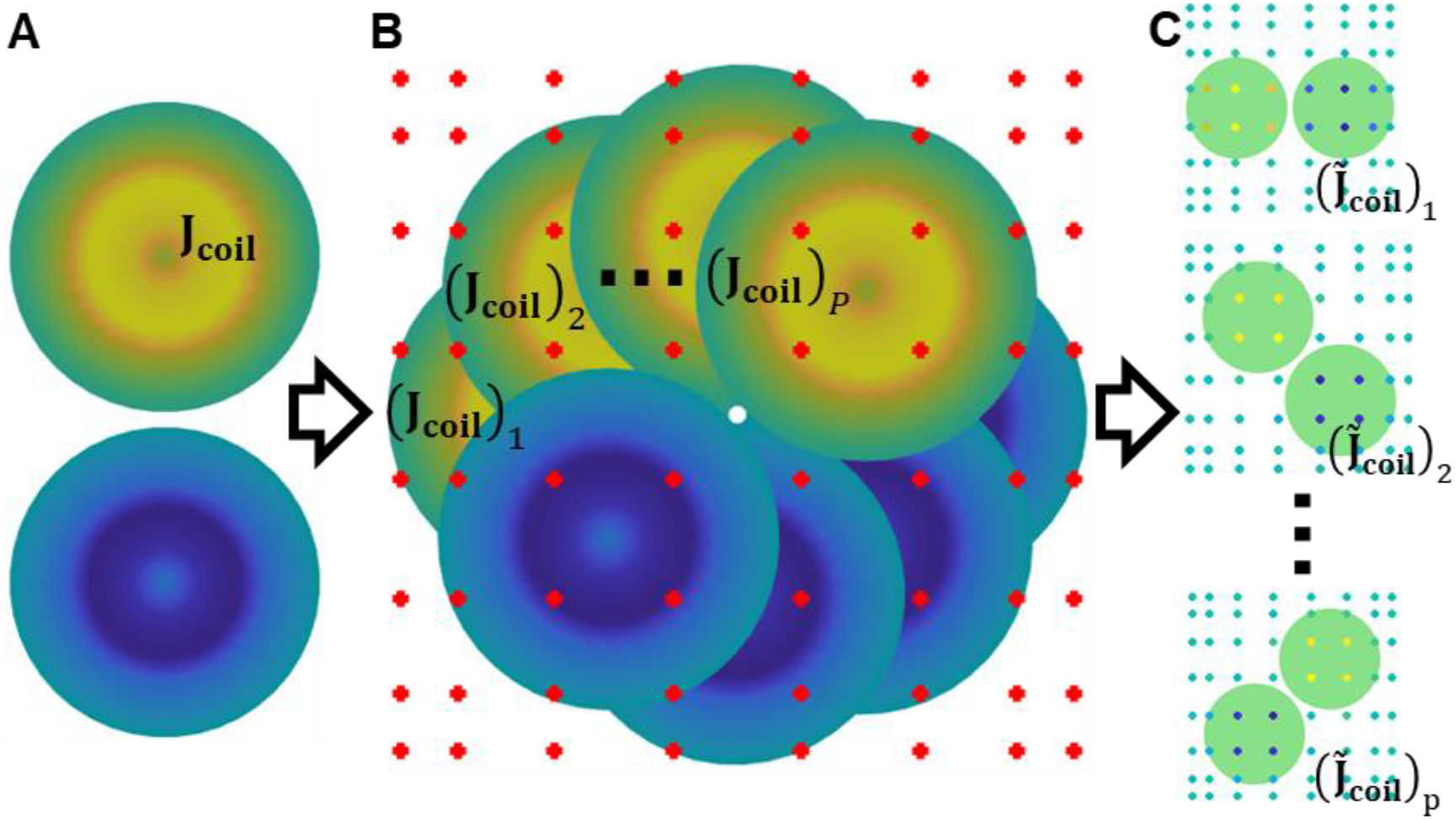
Auxiliary dipole method workflow. (A) TMS coil dipole model. (B) Model representing different TMS coil orientations along with dipoles placed on a grid of Gauss-Legendre nodes. (C) Dipole weights for individual coil orientations: warm and cold colors represent positive and negative values, respectively.

Since Ω_box_ is outside the head, which is a source-free region of space, the E-field can be interpolated as a polynomial with error decreasing exponentially with increasing degree of the polynomial[40]. The E-field samples 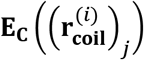 are determined from a polynomial fit of the Efield defined from auxiliary samples in Ω_box_.

Specifically, 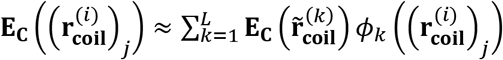, where positions 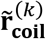 are chosen from a *N_x_× N_y_ × N_z_* tensor product of 1D Gauss-Legendre quadrature nodes in Ω_box_, and are chosen as their corresponding Lagrange polynomials. This choice of points finds a polynomial interpolant that minimizes L^2^ error in a subspace of polynomials with basis consisting of all monomials *x^i^y^j^z^k^*, where 0 ≤ *i* ≤ (*N_x_* – 1), 0 ≤ *j* ≤ (*N_y_* – 1), and 0 ≤ *i* ≤ (*N_z_* – 1) [49], thereby, resulting in a polynomial fit whose error decreases exponentially with increasing number of points. This method is agnostic to the choice of interpolant, as such, a reduced polynomial basis [50] and other spectrally accurate quadrature points [40], or a cylindrical bounding box along with a hybrid expansions of cardinal and Lagrange polynomial basis [49] could potentially be used to generate interpolants that achieve greater accuracy with less points.

Plugging in our interpolant into Eq.(7) results in the following:

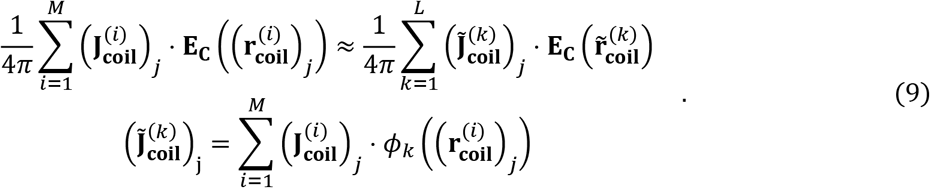

In Eq. (9), all of the coil orientations share the same dipole locations enabling the batch evaluation of **E_avg_** (Fig. 3C). Furthermore, Eq. (9) only incorporates information about the Lagrange interpolating polynomial functions; as such, it remains valid for magnetic dipole models without modification.

### Coil modeling

The 2712 magnetic dipole model of the Magstim 70 mm figure-8 coil (P/N 9790) [51] included in SimNIBS simulated for dI/dt = 1 A/μs was used for all accuracy comparisons (Fig. 1A). In its native space this coil is flat on the x-y plane, centered about the z axis. To keep computation times as small as possible, for all SimNIBS timing comparisons we used the precomputed magnetic vector potentials in lieu of the coil from the model package (file: ‘Magstim_70mm_Fig8.nii.gz’).

To speed-up computation further, we used the auxiliary dipole approximations of the coil to model each a distinct coil orientation. The dipole locations are the same across different auxiliary coil models, and coil models only differ in terms of their dipole weights. As a result, we can rapidly determine E-fields for many coil orientations each time using the same E-field samples outside of the head. In particular, we generated 360 models by rotating the coil dipole model around the z axis (0° to 359° in steps of 1°). To test the convergence of ADM with respect to increasing number of dipole locations, we used the SimNIBS Ernie example model and evaluated auxiliary coil dipole models consisting of all possible combinations of *N_x_* = *N_y_* varying from 1 to 29, and *N_Z_* varying from 1 to 3. For all other simulations, the total number of auxiliary dipoles is 578 and grid parameters were chosen as *N_x_* = *N_y_* = 17 and *N_z_* = 2.

### Head modeling

For numerical validation of the method we employed the SimNIBS example spherical shell and Ernie MRI-derived head model meshes. The spherical head mesh consists of a homogenous compartment with radius of 9.5 cm and conductivity of 0.33 S/m. The SimNIBS v3.1 Ernie head model mesh [18] consists of five homogenous compartments including white matter, gray matter, cerebrospinal fluid, skull, and skin. The scalp, skull, and cerebrospinal fluid conductivities were set to the default isotropic values of 0.465, 0.010, and 1.654 S/m, respectively. The gray and white matter compartments were assigned anisotropic conductivities to account for the fibered tissue structure. This was accomplished within SimNIBS by co-registering the diffusion-weighted imaging data employing a volume normalization of the brain conductivity tensor, where the geometric mean of the tensor eigenvalues was set to the default isotropic conductivities of 0.275 and 0.126 S/m for gray and white matter, respectively.

Additional MRI-based head models—M1, M2, M3, and M4—were borrowed from one of our experimental studies involving E-field-based dosing [37], corresponding to subject ids S262, S263, S266, and S269, respectively. Briefly, the models were constructed using structural T1-weighted (echoplanar sequence: Voxel Size= 1 mm^3^, TR= 7.148 ms, TE = 2.704 ms, Flip Angle = 12°, FOV = 256 mm^2^, Bandwidth = 127.8 Hz/pixel), T2-weighted (echo-planar sequence with fat saturation: Voxel Size = 0.9375 × 0.9375 × 2.0 mm^3^, TR = 4 s, TE = 77.23 ms, Flip Angle = 111°, FOV = 240 mm2, Bandwidth = 129.1 Hz/pixel), and diffusion-weighted scans (Single-shot echo-planar: Voxel Size = 2 mm^3^, TR = 17 s, TE = 91.4 ms, Flip Angle = 90°, FOV = 256 mm^2^, Bandwidth = 127.8 Hz/pixel, Matrix size = 1282, B-value = 2000 s/mm^2^, Diffusion directions = 26). Each model was first generated employing the co-registered T1-and T2-weighted MRI data to model major head tissues (scalp, skull, cerebrospinal fluid, and gray and white brain matter) represented as tetrahedral mesh elements. The number of tetrahedral mesh elements and nodes ranged 0.679–0.699 million and 3.82–3.92 million, respectively. Mesh conductivities were assigned following the same procedure as for the SimNIBS Ernie model.

### ROI and grid generation

For the spherical head, we considered a 1 cm diameter ROI centered 1.5 cm directly below the apex of the sphere as shown in Fig. 1C. A number of coil placements were chosen on a 2 cm diameter grid of 3.7-4.2 mm spaced positions and centered 4 mm above the apex of the sphere (Fig. 1C). For each coil placement, the coil was oriented tangentially to the sphere surface and different orientations where chosen by counterclockwise angular displacements of 0°, 45°,90°, and 135° relative to 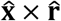. In other words, candidate orientations 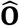 are 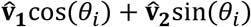, where *θ_i_*, is either 0°, 45°,90°, and 135°, 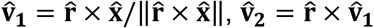, and 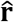 is the unit vector in the radial direction.

For the Ernie head model, the ROI was chosen as the gray matter contained in a 1 cm diameter cubic region centered about location (−55.1, −18.0, 96.4) as shown in Fig. 1D,E. Coil placements were chosen by extracting mesh nodes on the scalp that were within a 2 cm diameter of the point on the scalp closest to the centroid of the brain ROI and then projecting the nodes 4 mm outward in direction normal to the scalp surface. For each of the 632 coil placements (Fig. 1D), the coil was placed tangent to the scalp and different orientations where chosen by counterclockwise angular displacements of 0°, 45°, 90°, and 135° relative to 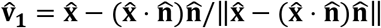, where 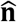 is the unit normal on the scalp directly below the center of the coil.

For each of the four additional models (M1-M4), four ROIs of various sizes where generated using the SimNIBS TMS coil placement optimization functionality. Within subject, the center of each ROI was the same and was derived from fMRI-measured brain activity [37]. The smallest ROI consisted of a single tetrahedron and had a mean effective diameter (the diameter of a sphere of the same volume) of 1.0 mm across subjects. The three larger ROIs were defined to have effective diameters of 10, 20, and 40 mm. SimNIBS was used to generate coil center positions on 3 cm diameter scalp region centered about the scalp position closest to the ROI’s center of mass [37]. Each coil position was chosen to be approximately 1 mm apart and scalp positions where extruded outward by an amount dependent on the individual hair thickness: 3.26, 2.03, 5.55, and 2.78 mm for models M1–M4, respectively [37]. The total number of coil positions was 697, 689, 689, and 697 for models M1–M4, respectively. For each position, we used the direct approach with SimNIBS and the reciprocity-based ADM to generate, respectively, 72 and 360 coil orientations tangent to the scalp by rotating the coil in 5° and 1° intervals about the scalp normal direction. The coarser discretization of the search space for SimNIBS was chosen to make the optimization computationally tractable.

### Visualizations

In this work, all visualizations of anatomical details of the volume conductor models, TMS coil setups and optimizations, and E-field distributions are generated with SCIRun (v4.7, R45838; [30], Figures 1 and 9 and parts of Figures 10 and 11) and MATLAB R2019 (Figures 3–7, 10, and 11)[52].

### Error metrics

For each optimization setup we consider a normalized absolute error metric, which is normalized to the maximum observed 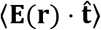,

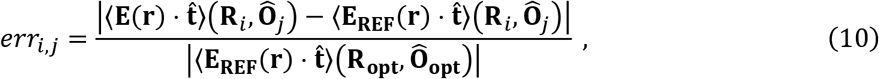

where 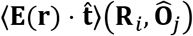 and 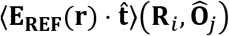 are the average E-field components in direction 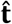 computed by one of our proposed methods and a reference method, respectively, for the *i*^th^ candidate coil position and *j*^th^ orientation. Additionally, we compare the E-field magnitude estimate 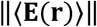 to 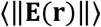 normalized to the maximum observed 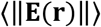,

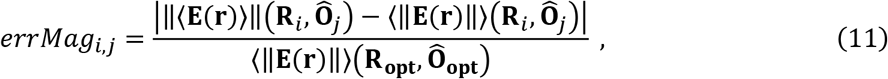

### Coil placement uncertainty quantification

Using the reciprocity technique, we can also compute uncertainty in the E-field resulting from coil placement uncertainty. First, we use the reciprocity method to construct maps indicating the Efield induced for all possible coil positions and orientations on the considered region of the scalp. Second, this map is used to extract uncertainty of the E-field induced during TMS resulting from uncertainty in coil placement.

Uncertainty in coil placement is modeled as a probability distribution describing the likelihood of the coil being placed with a given orientation and location on the scalp. Here we assume that the coil position and orientation are uniformly distributed random variables with distributions centered at the nominal scalp position **R_μ_** and coil orientation 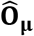. The support of the distribution comprises the scalp region within a geodesic distance *R*_Δ_ from **R_μ_** and all coil orientation deviations up to *θ*_Δ_ from 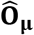. Specifically, the support is

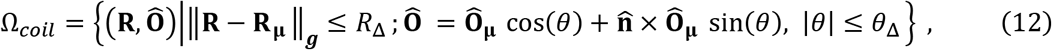

where ||**R – R_μ_**||_*g*_ is the geodesic distance along the scalp between **R** and **R_μ_**, and parameters *R*_Δ_ and *θ*_Δ_ are radii of the positional and angular support of the distribution. The geodesic distance on the scalp is computed using the heat method described in [53]. The values of *R*_Δ_ and *θ*_Δ_ are specific to the TMS methods used and should be based on available data [24–26]. Other distributions (e.g., Gaussians) could also be used to discretize the coil position and orientation uncertainty whenever appropriate.

## Results

### Validation

For the spherical head model, we compare the average E-field in the ROI predicted by our in-house direct FEM, reciprocity, and ADM with the analytically computed value. The average Efield computed analytically for each coil placement is shown in Fig. 4. As expected, the maximum occurs when the coil is placed directly above the ROI and oriented to induce a maximum primary E-field along 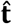. Fig. 4 also shows the errors normalized to the maximum E-field across all simulations. The maximum error obtained for the direct FEM, reciprocity, and ADM are 0.06%, 0.08%, and 0.46%, respectively. There is a slightly reduced accuracy in the ADM relative to the direct FEM and reciprocity approaches. However, the errors are still very low with the ADM compared to modeling and numerical errors that can exceed 5% in TMS E-field simulations[33,43] This indicates that 578 dipoles are sufficient to accurately represent the coil fields for all the test cases. These simulations where also run in SimNIBS and we observed maximum error for SimNIBS simulations of 0.16%. Additional SimNIBS error figures are given in the supplemental material.

**Figure 4.**
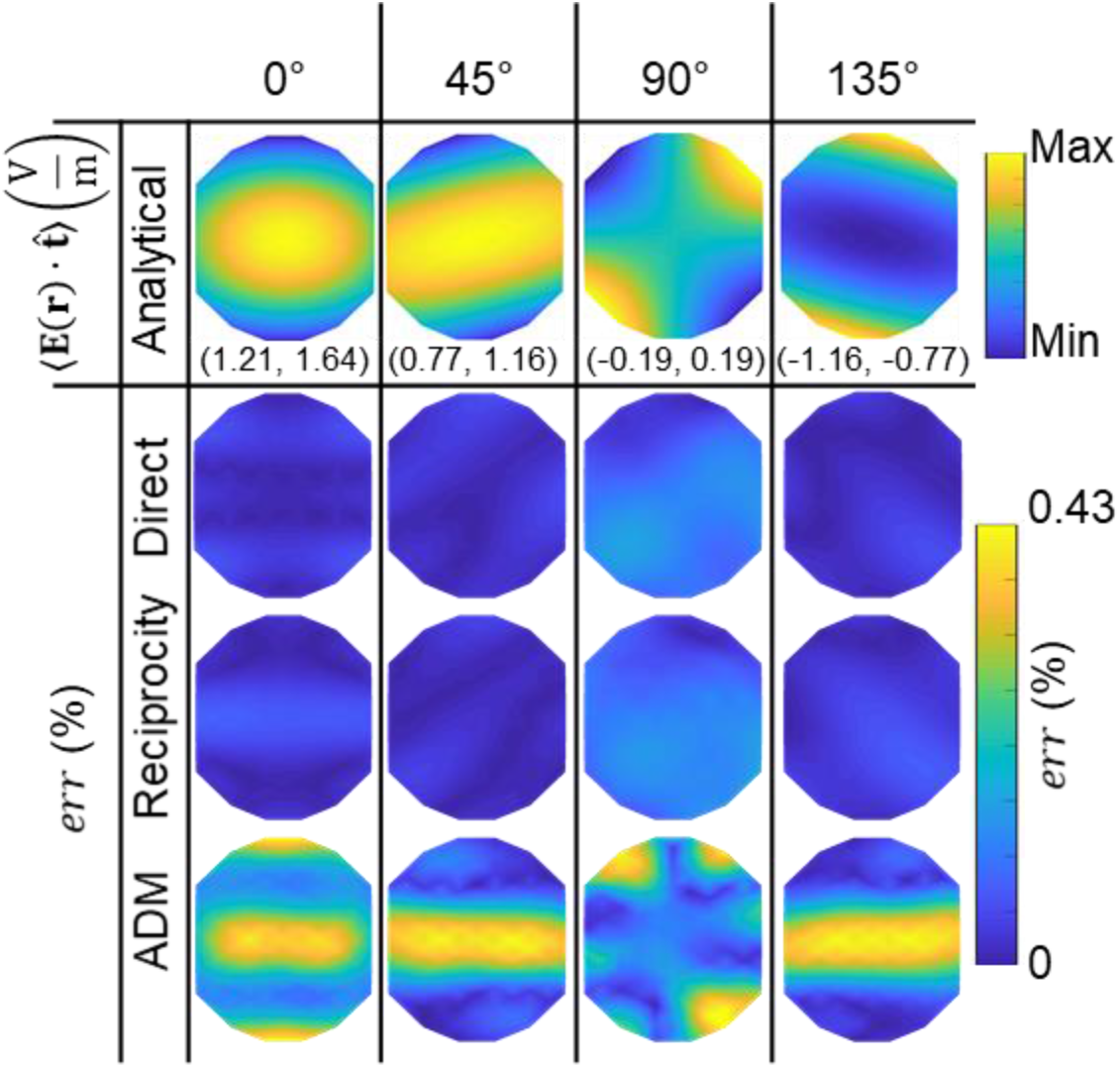
Validation of the computational methods in sphere head model. The 1^st^ row shows, as a reference, the analytically computed average ROI E-field in direction t (oriented horizontally) across TMS coil positions on the scalp. The scalp area spanned by the coil positions has a diameter of 2 cm. Four coil orientations are considered, from left to right: 0°, 45°, 90°, and 135°. The range of observed E-field component values are given in parenthesis below each figure in V/m. The 2^nd^, 3^rd^, and 4^th^ row contain the corresponding absolute error relative to the analytical solution, *err*, for the direct, reciprocity, and ADM approaches, respectively.

For the Ernie head model, we compare the average E-field in the ROI predicted by the in-house direct FEM, reciprocity, and ADM. The value of 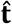 is chosen (as described above) as (−0.57,−0.72,−0.39) by evaluating the E-field for coil orientations varying from 0° to 359° in 5° intervals at each of the coil positions. The average E-field computed by the direct FEM for each coil position is shown in Fig. 5. A maximum E-field is obtained when the coil is placed 4.5 mm away from the point on the grid closest to the ROI center of mass. The normalized absolute error of the different approaches, when using direct FEM as reference, is shown in Fig. 5. All approaches agree with the direct FEM and have relative absolute errors below 0.13%. The maximum normalized absolute errors across all simulations of ADM relative to the reciprocity approach is shown in Fig. 6. ADM converges to the reciprocity results exponentially with increasing number of dipoles; an error below 0.5% is observed for the configuration of 578 dipoles used throughout this paper. Additional comparisons with SimNIBS results given in the supplemental material have a maximum relative error below 0.11 %.

**Figure 5.**
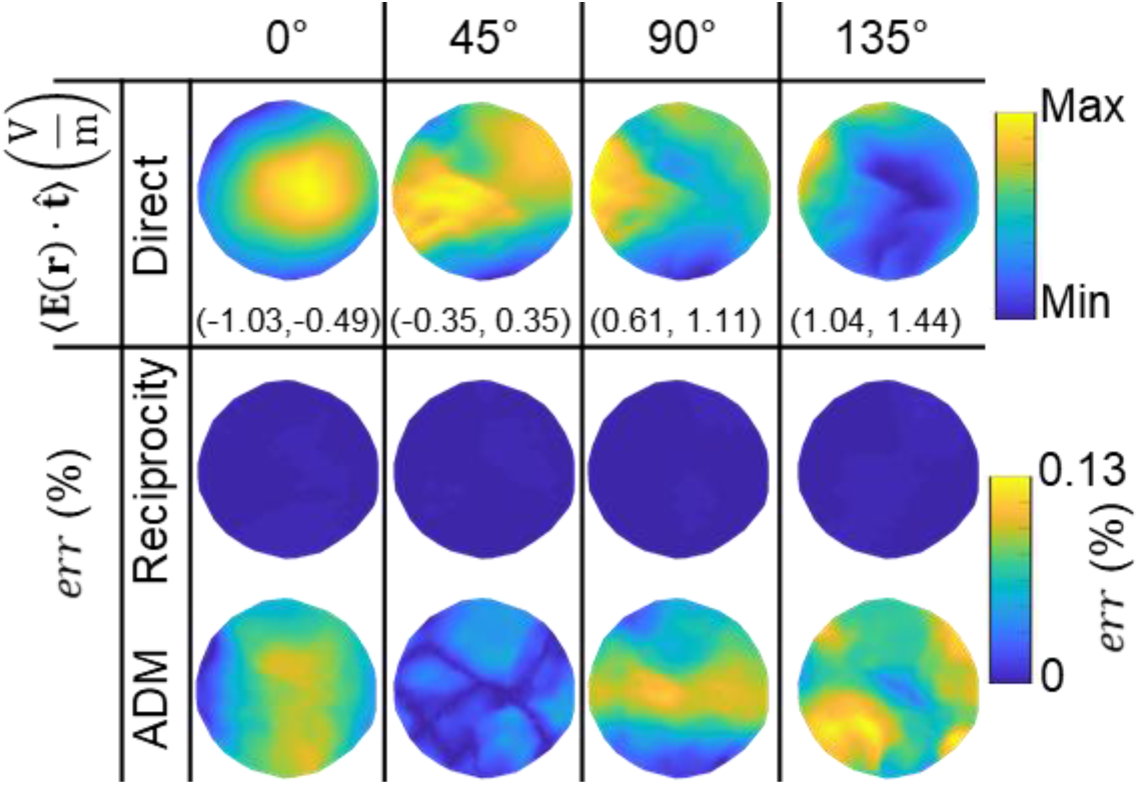
Validation in the Ernie head model, analogous to Fig. 4. The 1^st^ row shows, as a reference, the directly computed average ROI E-field in direction 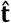 (oriented horizontally) across TMS coil positions on the scalp. The scalp area spanned by the coil positions has a diameter of 2 cm. Four coil orientations are considered, from left to right: 0°, 45°, 90°, and 135°. The range of observed E-field component values are given in parenthesis below each figure in V/m. The 2^nd^ and 3^rd^ row contain the corresponding error relative to the direct approach, *err*, for the reciprocity and ADM approaches, respectively. The direct approach is used as a reference since there is no analytical solution for anatomically-detailed MRI-based head models.

For the four models (M1–M4) we compared metric 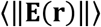 to 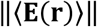 for ROIs of varying sizes. Fig. 7 shows the maximum magnitude error obtained by comparing ADM to compute 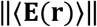 relative to using direct FEM to compute 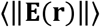 for ROIs of varying sizes. The maximum error increases with increasing ROI size. Considering the spatial resolution of TMS, the diameter of a typical ROI is below 1 cm diameter, and for these sizes the errors are below 0.5 %. For larger ROIs with diameters between 1 cm and 2 cm the error is still below 2.0 %, commensurate with the numerical error of good TMS E-field solvers [33]. These results indicate that 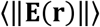 can be approximated by 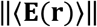 in many practical TMS settings.

**Figure 6.**
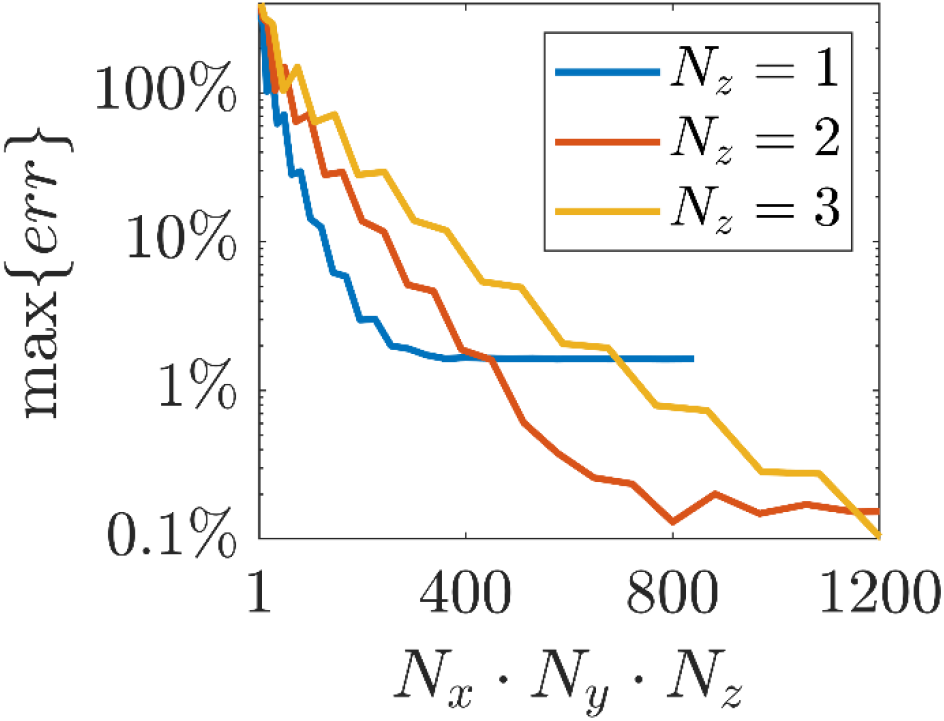
Maximum normalized absolute error for ADM relative to the reciprocity results as a function of number of ADM auxiliary dipoles *N_x_* · *N_y_* · *N_z_*.

**Figure 7.**
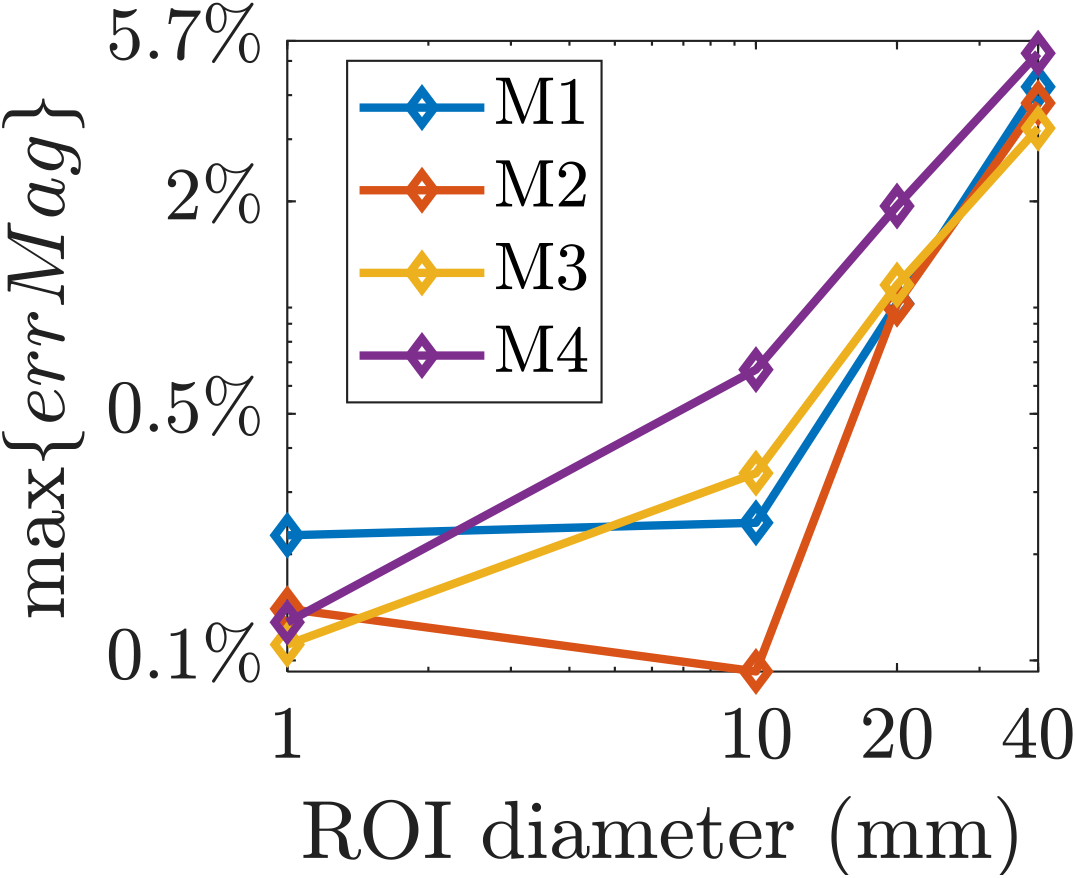
Maximum E-field magnitude estimate error (Eq. (9)) using ADM compared to the reciprocity results as a function of ROI diameter.

### Computational runtime and memory

Coil placement optimizations were run each time using a different number of candidate designs. The optimizations were run using a high performance computation system that has an Intel(R) Xeon(R) Gold 6154 CPU with a 3.0 GHz 36 core processor and 768 GB of memory as well as a conventional laptop that has an Intel(R) Core(TM) i7-4600U CPU with a 2.10 GHz dual-core processor and 12 GB of memory. For each model (M1-M4) we ran the SimNIBS coil position and orientation optimization procedure to maximize 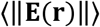 and ADM to maximize 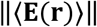. Fig. 8 shows computational times required as a function of total candidate coil positions and orientations.

**Figure 8.**
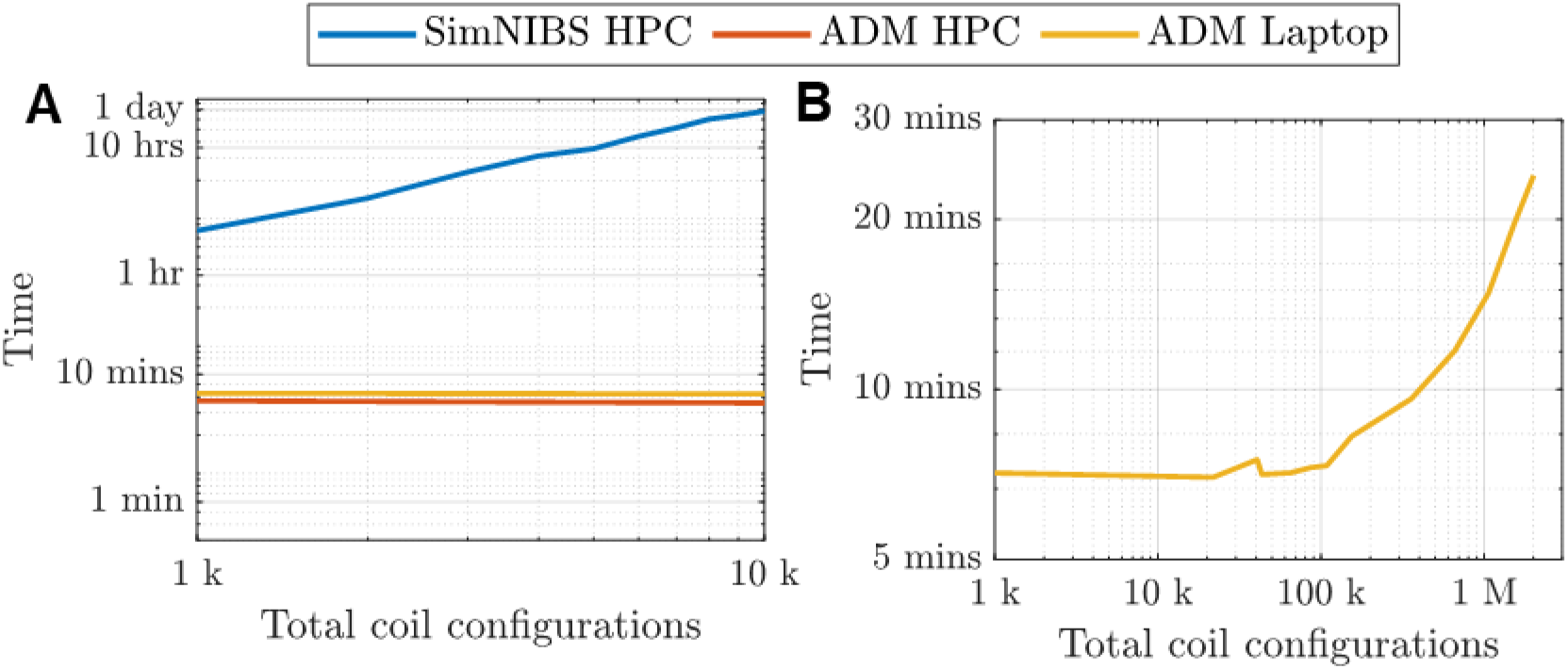
CPU runtime versus number of total coil configurations. (A) Results for SimNIBS and ADM run on a high performance computation system (HPC) and a laptop (ADM only). (B) Extended results for ADM. CPU runtimes are averaged across models M1–M4.

**Figure 9.**
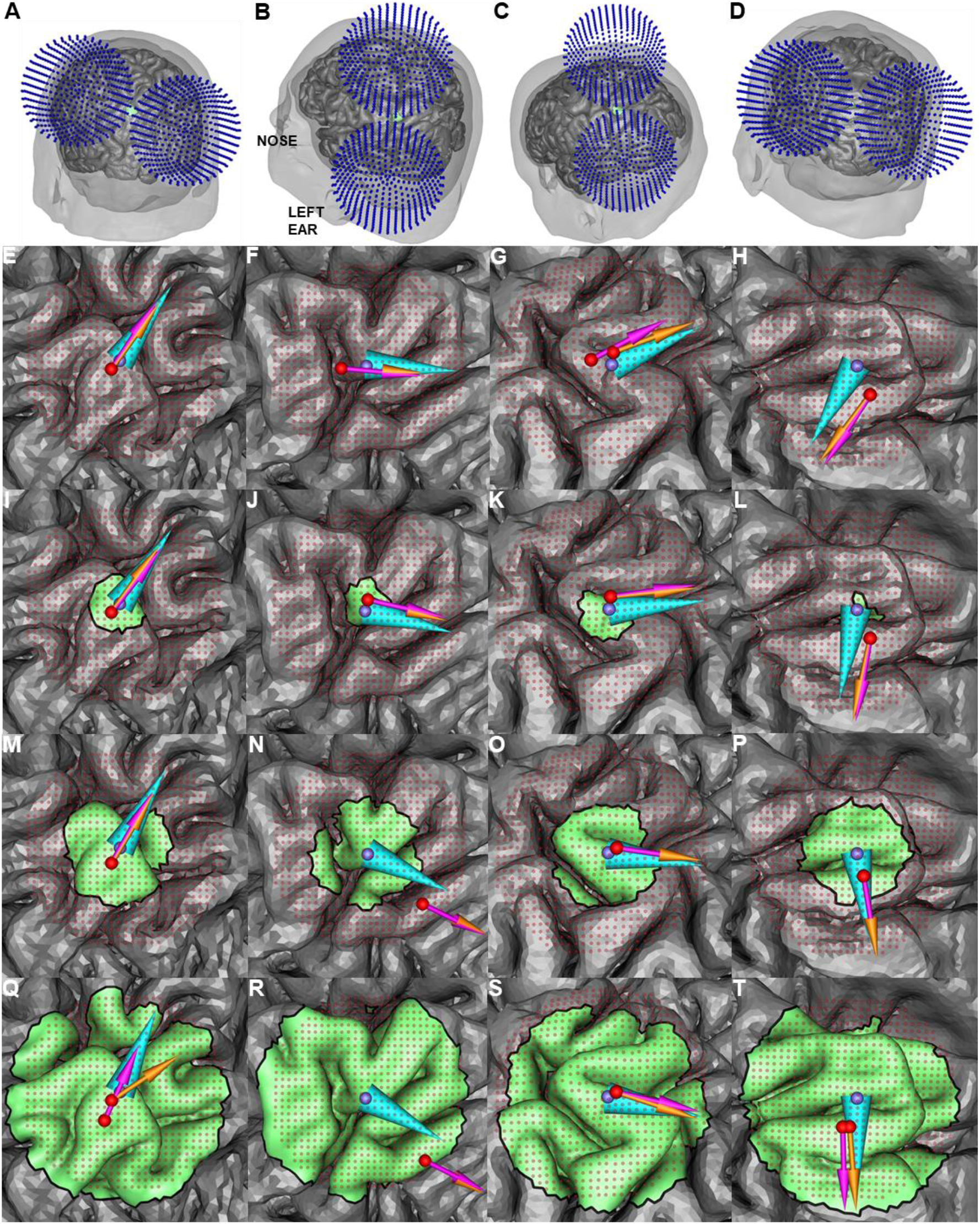
Optimal coil placement results for ROIs of various size in models M1–M4 (left to right columns). (A)-(D): Illustration of the ADM-optimized coil placement for the 10 mm diameter ROI in models M1–M4, respectively. (E)-(T): SimNIBS and ADM optimized position and orientation are represented by purple and orange arrow, respectively, 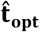 and the ROI center of mass are represented by cyan cone and purple sphere, respectively. Rows, top to bottom, show results for increasing ROI diameter of 1, 10, 20, and 40 mm. In most cases both optimization methods result in the same or similar coil position and orientation.

On the high performance computation system, the SimNIBS coil placement optimization required on average 7 seconds per candidate configuration, whereas our approach required an average total time of 7 minutes and 15 seconds regardless of the number of candidate configurations. As such, a SimNIBS optimization with 10000 candidate coil placements takes 20 hours, whereas ADM is 165 times faster, taking a little over 7 minutes. The memory required to run the SimNIBS optimization ranged 11–11.5 GB, whereas with ADM it was only 6.3–6.5 GB. Note that the memory requirements of SimNIBS can be lowered to 3.5 GB by using their iterative solver for the optimization [43], which, however, takes significantly longer. We decided not to use the SimNIBS iterative solver because many current laptop computers have at least 16 GB of memory and, thus, memory requirements do not impede coil position and orientation optimization. ADM computational time starts to increase beyond 100 thousand coil placements (Fig. 8B). The total time to run 1 million coil placements was 15 minutes, whereas assuming 7 seconds per simulation, the SimNIBS direct approach would have taken approximately 8 days.

### Example application of coil position and orientation optimization

Fig. 9 shows results for models M1–M4 using SimNIBS to maximize 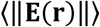 and ADM to maximize 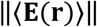. Except for model M3 and its 1 mm (one tetrahedron) diameter ROI, and 40 mm diameter ROI results, all of the coil placement optimizations resulted in the same position and differ in orientation by less than 2°. For model M3 and the 1 mm diameter ROI, the optimum coil position computed by our in-house direct method and the ADM approach are identical (supplemental Fig. S6) indicating that the 3.3 mm distance between the optimum of SimNIBS and ADM are likely due to numerical errors. For ROIs with 40 mm diameter, the discrepancy between the ADM and SimNIBS optimum TMS coil positions is likely due to inaccuracies in the approximation of the E-field magnitude using ADM. For regions with diameter below 40 mm, the orientation differences are likely because ADM was run with an angular step size of 1°, whereas the SimNIBS simulations were run with an angular step size of 5° (due to long computation times). As such, the observed differences in coil orientation between ADM and SimNIBS are below the resolution of the SimNIBS optimization.

Fig. 9 shows the E-field orientation associated with maximum E-field magnitude, 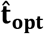. As is generally the case for TMS, the strongest E-field is approximately tangential to the scalp surface {Ilmoniemi, 2019 #7}. Relative to the cortical gyri, 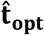 is typically nearly perpendicular to the nearby sulcal walls. This is consistent with other modeling and experimental results [14,18,19]. Whenever there are multiple bends in the ROI, it is difficult to identify the precise normal to the sulcal wall. Our framework helps to disambiguate a specific coil orientation that should be used, although in some cases a range of orientations may yield similar E-field strengths at the target.

In a conventional neuronavigated coil placement protocol, the coil would be placed directly above the center of mass (**R_CM_**). We compare the optimum coil placements to a placement where the coil is placed above **R_CM_** and oriented along normal to the nearest sulcal wall to the ROI center of mass (Fig. 9). Fig. 10 compares the coil placement above the center of mass and the SimNIBS and ADM optimization approaches. The average E-field magnitude in the ROI obtained by placing the coil at **R_CM_** can be up to 6.0% lower than that for the E-field guided methods (Fig. 10A-B). The E-fields achieved by each optimum coil placement strategy are almost identical, differing by less than 0.3% (Fig. 10C). The coil position directly above the ROI center of mass and the optimization methods can vary as much as 13.8 mm (Fig. 10D-E), whereas this difference is less than 3.2 mm between ADM and SimNIBS (Fig. 10F). Thus, the ADM method to optimize 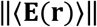 results in nearly the same E-field delivery and coil location as using SimNIBS to optimize 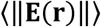, and both approaches are superior to the coil placement directly above the ROI center.

**Figure 10.**
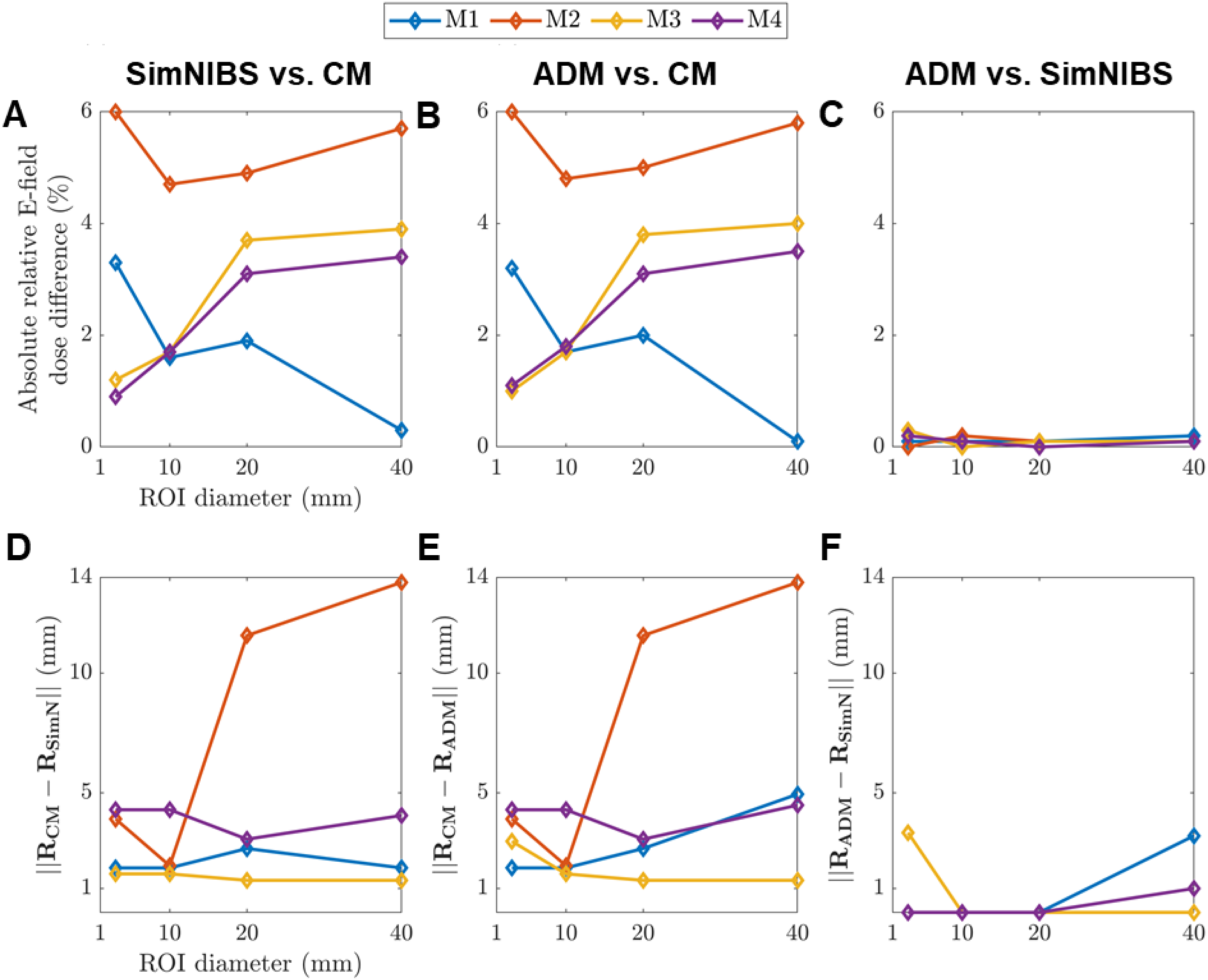
Comparisons of induced average E-field magnitude (A-C) and coil position (D-F) for ROIs of various sizes across different coil positioning strategies: (A,D) SimNIBS optimization versus a placement over ROI center of mass; (B,E) ADM optimization versus ROI center of mass placement; (C,F) ADM versus SimNIBS. Additional coil optimization comparisons are given in Tables S1 and S2 in the supplemental material.

### Coil placement uncertainty quantification

As an example, we computed the uncertainty in 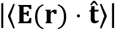 for the 10 mm diameter ROI in head model M2 resulting from coil position and orientation uncertainty. Fig. 11 shows results for various levels of coil position uncertainty. For fixed *R_Δ_*, the minimum of the standard deviation is where the expected value of 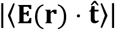 is highest. The standard deviation for each position increased with *R_Δ_*. For fixed *R_Δ_*, the expected value decreased monotonically away from its optimum, and this decrease accelerated with increasing. Fig. 12 plots the 90% confidence interval and expected value for 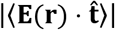 for various levels of position and orientation uncertainty. The expected value of 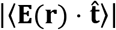 decreased with increasing. The range of the 90% confidence interval of 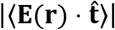 nearly doubled if the coil was placed 5 mm away from its optimal position (Fig. 12A-E). Furthermore, coil orientation uncertainty did not contribute significantly to the total uncertainty (Fig. 12F). These results indicate that errors in identifying the optimum coil position both decrease effectiveness of TMS and also increase uncertainty in the TMS induced E-field due possible coil positioning errors. Furthermore, it is more critical to identify the optimum coil positioning whenever there are large coil positioning uncertainties associated with the TMS coil placement protocol.

**Figure 11.**
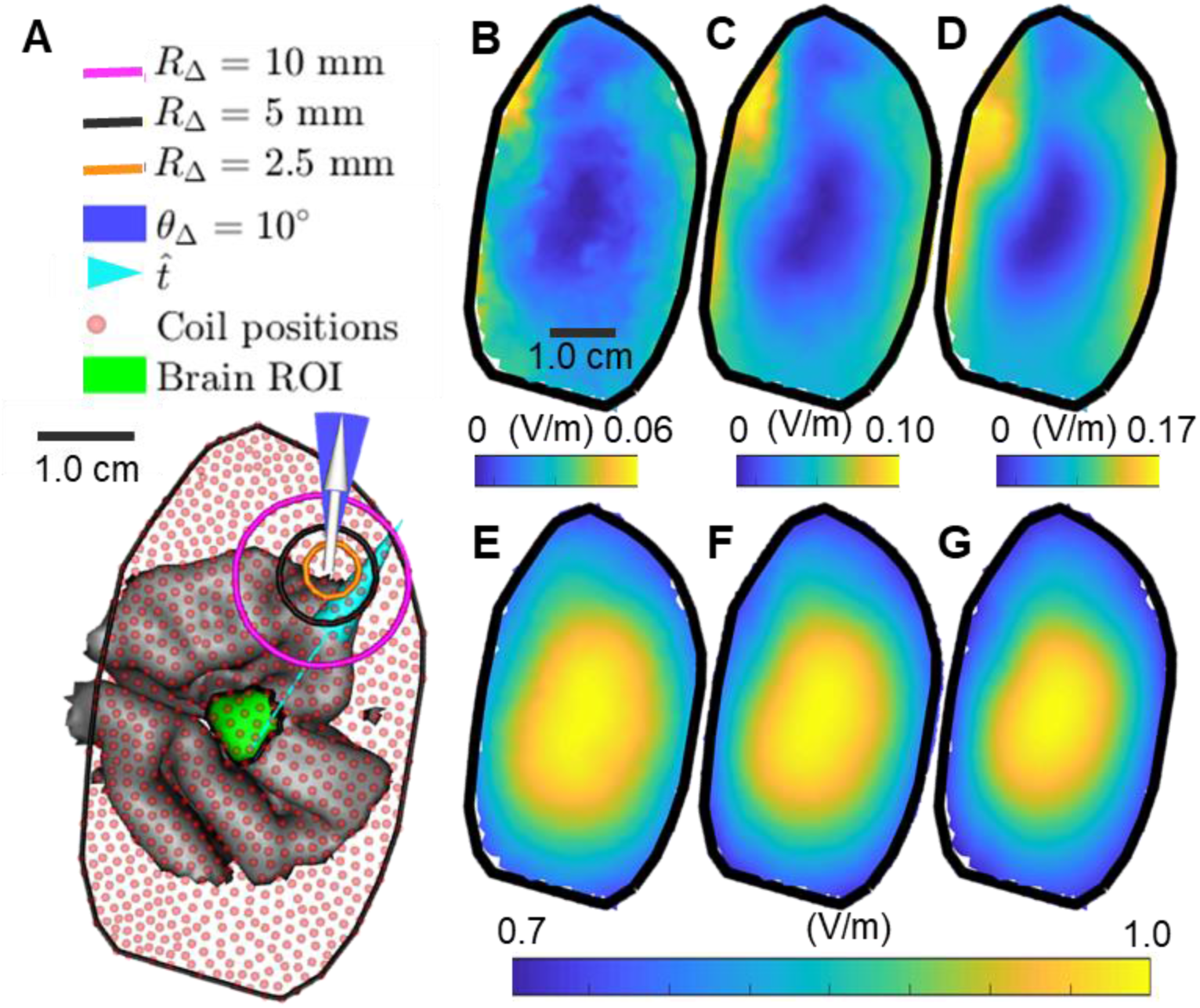
Coil position uncertainty results for model M2 and 10 mm diameter ROI. (A) Coil placements are chosen on the scalp above the brain ROI. The coil is oriented along the white vector and orientation uncertainty is always chosen as *θ*_Δ_ = 10°. The support of coil position uncertainty for *R_Δ_* = 2.5 mm, mm, *R_Δ_* = 5.0 mm, and *R*_Δ_, = 10 mm is marked by the orange, black, and magenta circles, respectively. The maximum of the expected value and standard deviation for the average E-field along 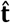 for each coil position is determined. (B-D) The standard deviation assuming a coil position uncertainty of (B) 2.5 mm, (C) 5 mm, and (D) 10 mm. (E-G) The expected value for the average E-field assuming a coil position uncertainty of *R*_Δ_ (E) 2.5 mm, (F) 5 mm, and (G) 10 mm. Results are normalized by the maximum expected average E-field over all coil positions and orientations.

**Figure 12.**
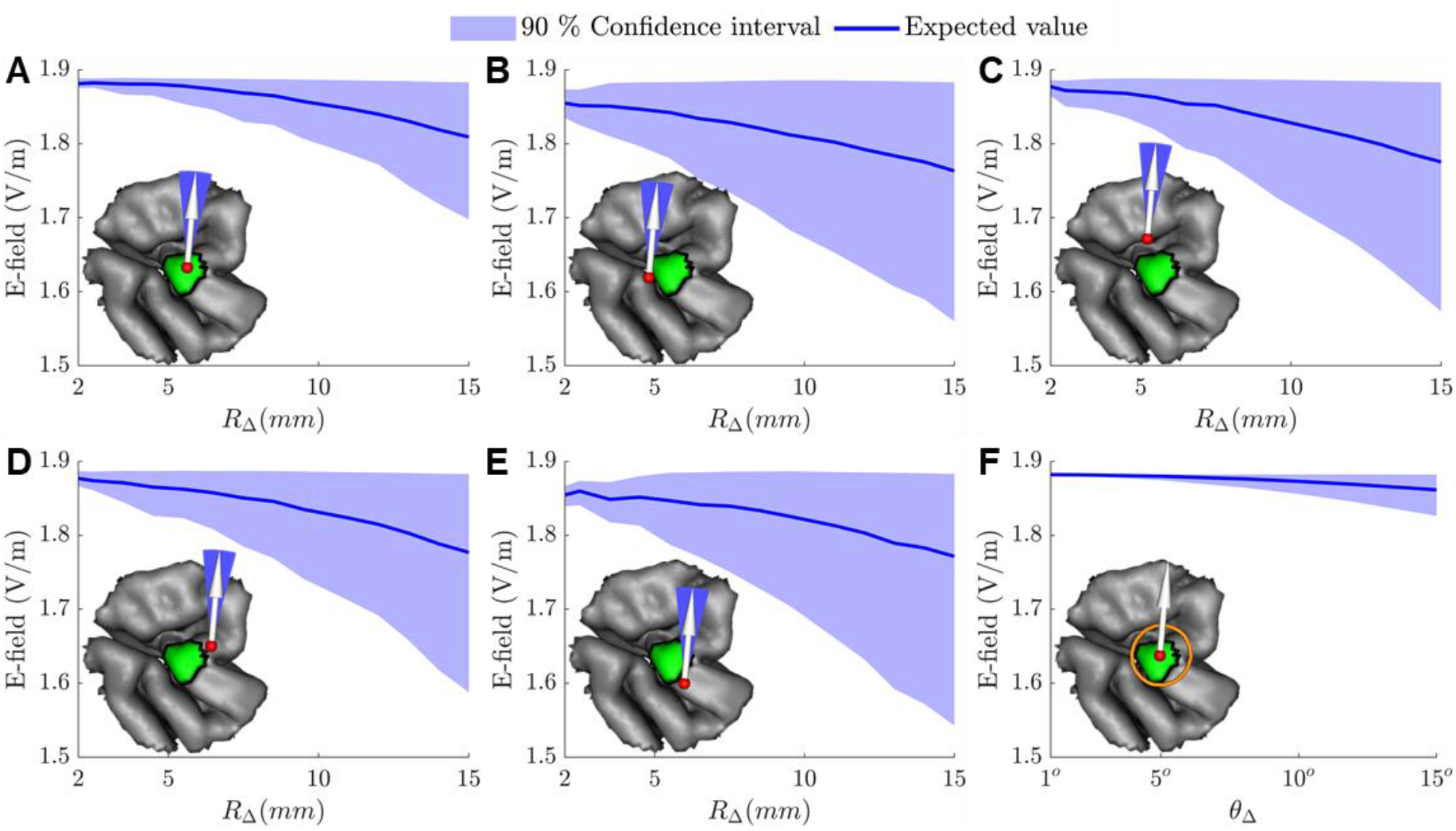
90% confidence region of marginal distributions and expected value of the average E-field along t in the ROI as a function of coil position and orientation uncertainty. (A-E) Results for *θ*_Δ_ = 10° and the coil positioned (A) centered or (B-E) 5 mm off-center relative to the ROI. (F) Results for *R*_Δ_ =5 mm and the coil centered above the ROI.

## Discussion and Conclusion

ADM, which uses auxiliary dipoles along with electromagnetic reciprocity, enables rapid extraction of optimal coil position and orientation to target the TMS E-field to specific brain regions. Furthermore, ADM enables the rapid quantification of uncertainty of TMS induced ROI E-fields resulting from uncertainty in the coil position and orientation. The validation results indicate that the average E-field along a given direction in an ROI can be computed with a numerical error below 1% using ADM with 578 dipoles. For ROIs with diameter below 2 cm, the average magnitude of the E-field can be estimated to an error below 2% by executing ADM only three times, corresponding to three orthogonal spatial directions of the E-field. As such, ADM can accurately determine the optimal coil placement based on maximization of either the E-field along a given direction or the total E-field magnitude for typical cortical ROIs. The optimal coil positions and orientations determined via ADM and the direct FEM approach for maximizing the average E-field are virtually identical for ROIs with diameter below 2 cm. However, ADM runs in under 15 minutes on a laptop to optimize the coil placement, whereas the direct FEM computations take over two days. Thus, ADM is an accurate and rapid method for E-field-informed coil placement that can be implemented on a standard laptop.

Supporting the value of coil placement optimization, the difference between the location on the scalp closest to the targeted ROI center of mass, which represents conventional neuronavigated targeting, and the E-field-informed optimum coil position is 1-14 mm. This is consistent with prior results using direct simulations that observed an average distance of 5.5 mm and as high as 12 mm for ROIs on the temporal brain region [14].

The ADM method can be used with a pre-specified optimal E-field direction, if such is known or hypothesized based on a directional specificity of the activation of the targeted neural elements [4]. When the objective is to maximize the E-field magnitude in the ROI, the optimal E-field direction is typically close to perpendicular relative to the sulcal wall, consistent with prior studies using the direct method [14,18,19]. However, the optimal coil orientation is ambiguous for ROIs containing highly curved sulcal walls, which means that a range of coil orientations can produce essentially the same E-field magnitude at the cortical target.

The quantification of E-field uncertainty shows how errors in coil positioning both decrease the E-field strength at the target and increase its variability. If there are significant coil positioning uncertainties associated with the TMS protocol, the importance of determining the optimum nominal coil position increases since it can prevent excessive variability in the E-field delivered to the target. Finally, the estimated E-field variation due to procedural uncertainty can enable statistical analysis involving the E-field dose.

## Conflict of interest declaration

The authors declare that there is no conflict of interest regarding the publication of this article.

## Acknowledgement

Research reported in this publication was supported by the National Institute of Mental Health and the National Institute on Aging of the National Institutes of Health under Award Numbers K99MH120046, RF1MH114268, RF1MH114253, and U01AG050618. The content of current research is solely the responsibility of the authors and does not necessarily represent the official views of the National Institutes of Health. We thank Dr. Boshuo Wang and Dr. Lari Koponen for providing helpful conversations and suggestions on this study.

## Supplemental Material

### Reciprocity method applied to magnetic dipole coil models

**Figure S1.**
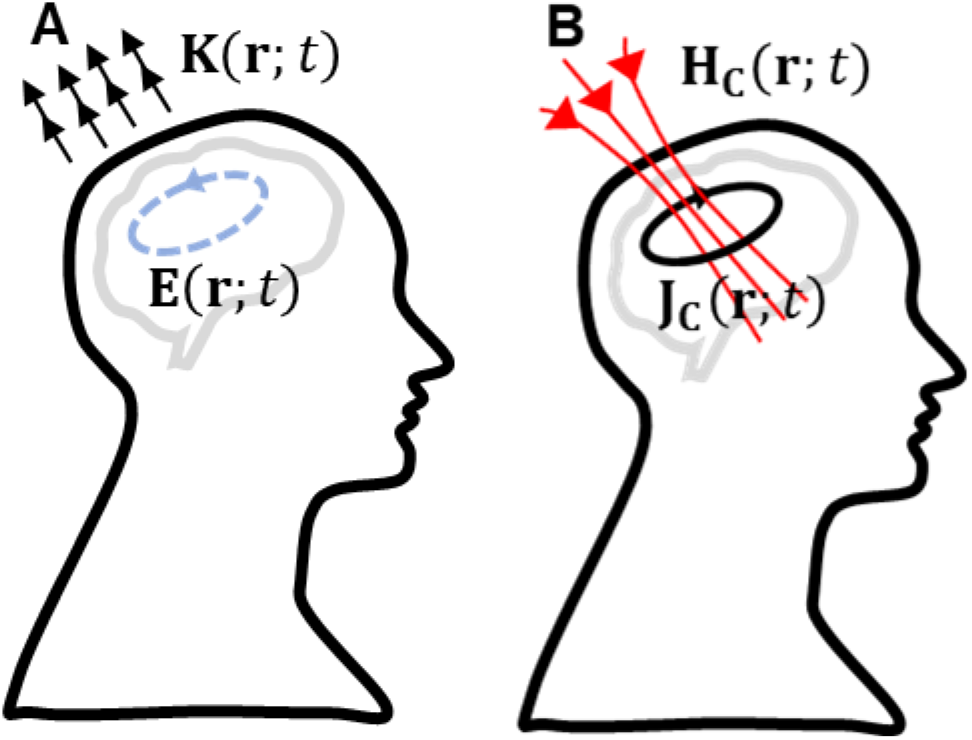
Alternative reciprocal scenarios: (A) Magnetic dipoles representing the TMS coil generate an E-field inside the cortex. (B) A brain current source generates a magnetic field (H-field) where the coil resides.

Here the reciprocity section of the main text is repeated with appropriate field related quantities replaced to allow the use of a magnetic dipole model of the TMS coil. In this case, electromagnetic reciprocity is an equivalence relationship between two scenarios (Fig. S1). In one scenario, the TMS coil, modeled as impressed magnetic currents **K**(**r**;*t*) = *p*’(*t*)**K**(**r**), generates an E-field **E**(**r**; *t*) = *p*’(*t*)**E**(**r**) inside the head (Fig. S1A). In the second scenario, a source current **J_c_**(**r**; *t*) = *p*(*t*)**J_c_**(**r**) inside the head generates an H-field **H_c_**(**r**; *t*) = *p*(<u>t</u>)**H_c_**(**r**) where the coil resides (Fig. S1B). Reciprocity dictates that the reaction integral between **E**(**r**;*t*) and **J_c_**(**r**;*t*) is equal to the reaction integral between **K**(**r**; *t*) and **H_c_**(**r**; *t*),

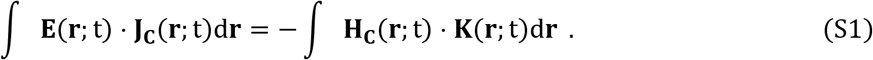

Here we choose **J_c_**(**r**;*t*) = 0 outside of the ROI and 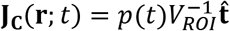 inside it. Reciprocity results in the following equation for the average E-field along 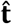 in the ROI

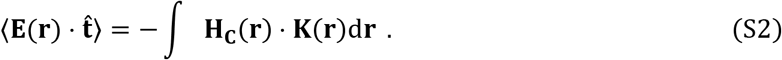

Note that the temporal variation has been factored out in this equation. In what follows, the temporal variation of all currents and E-fields is *p*(*t*) and *p*’(*t*), respectively, and is suppressed in the notation, as in the main text.

Computing 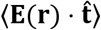 using Eq. (S2) requires the evaluation of **H_c_**(**r**) outside the conductive head. This is done by computing the primary H-field due to the cortical current **J_c_**(**r**) and secondary H-field due to conduction currents 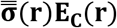, where 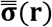 is the conductivity tensor inside the head,

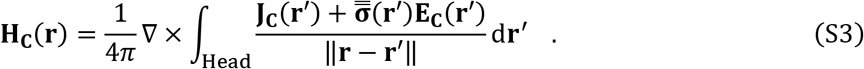

Finally, to estimate Eqs. (S2)–(S3) we use the same approach as described in the main text to estimate Eqs. (2)–(3). Specifically, Eqs. (5)–(6) become, respectively,

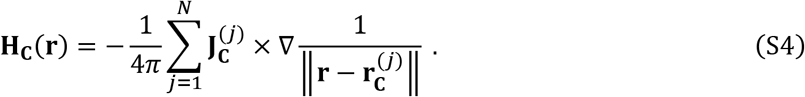

and

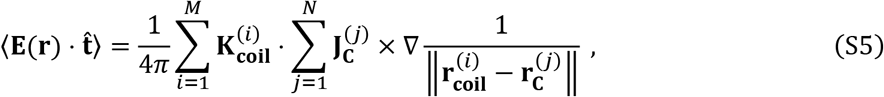

where magnetic dipoles 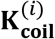 at locations 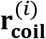 model the coil and *i* = 1,2, …,*M*.

### Validation comparison between simulations with SimNIBS and in-house FEM solver

Here additional validation results are given indicating that our in-house FEM implementation [1] produces E-field simulation results and error levels comparable to those of SimNIBS, a commonly used non-invasive brain stimulation simulation package [2,3]. Fig. S3 shows spherical head model results indicating that the SimNIBS errors are slightly higher than our direct FEM (see Fig. 3 in the main text). These small differences in accuracy are likely due to differences in the way that the primary E-field is computed. Fig. S4 compares the results obtained for the Ernie head model via our direct FEM with SimNIBS, indicating a maximum relative difference across all simulations of 0.11%. Fig. S5 compares results obtained for the four additional models (M1-M4) obtained via our direct FEM and SimNIBS. The agreement between the two sets of solutions increases with ROI size. This is likely because we compute the average the E-field on the ROI, which is less sensitive to numerical errors with increasing ROI size. In all cases, SimNIBS was in agreement with our direct FEM to a fraction of a percentage indicating that both codes result in virtually the same solution.

**Figure S3.**
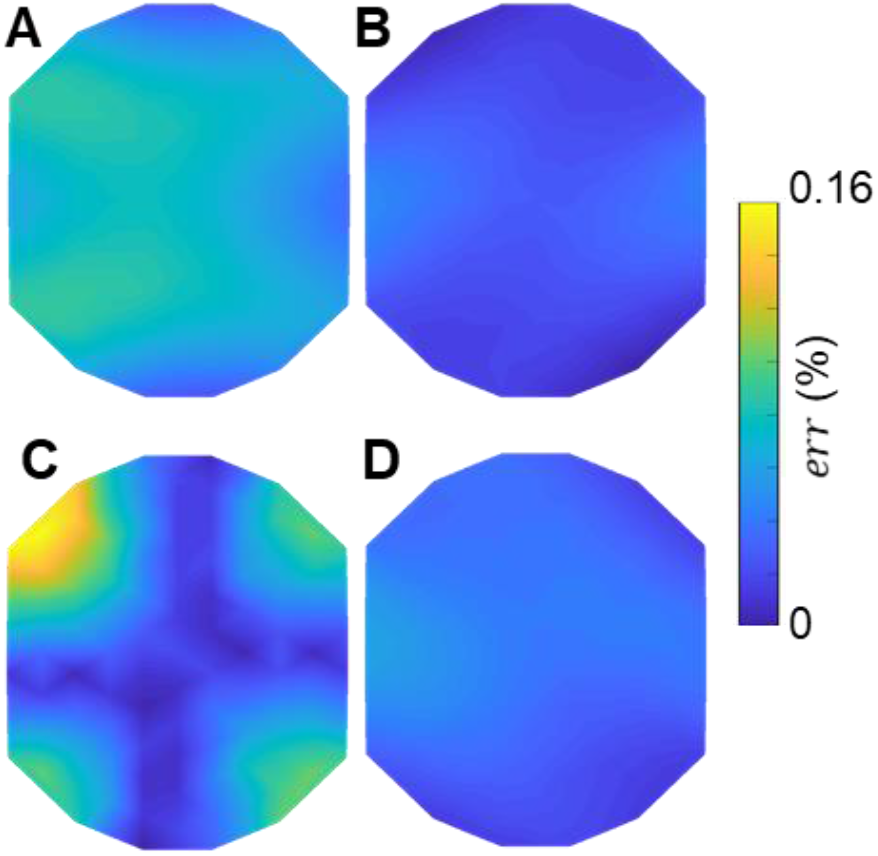
Error in SimNIBS E-field simulations relative to analytical solution in sphere head model, analogous to Fig. 3 in the main text. The plots show *err* observed at each scalp location for coil orientations of (A) 0°, (B) 45°, (C) 90°, and (D) 135°.

**Figure S4.**
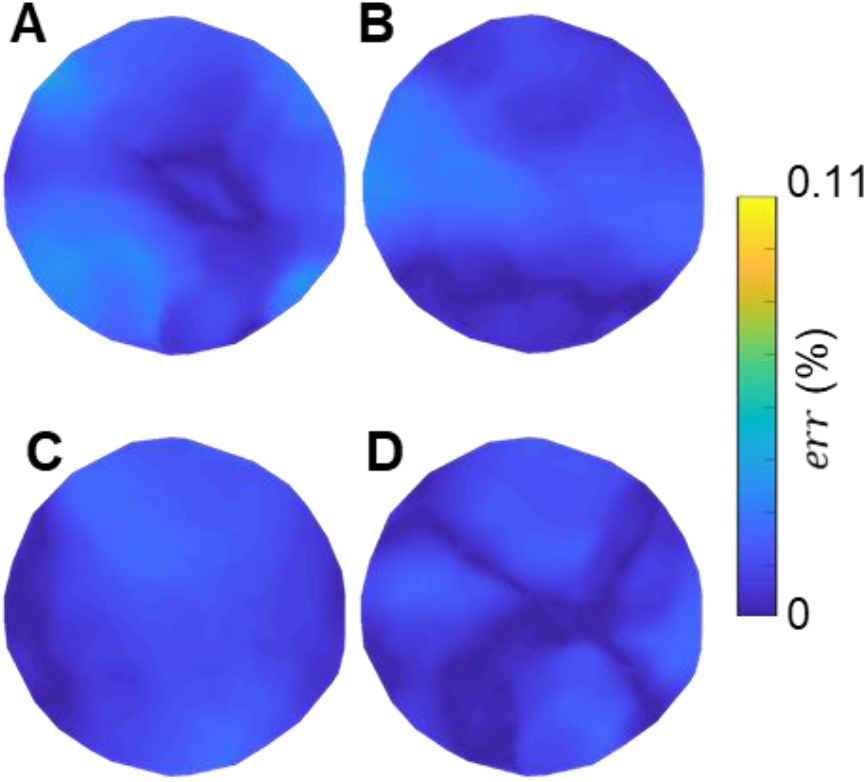
Difference in SimNIBS simulation results for Ernie head model relative to our direct FEM results, analogous to Fig. 4 in the main text. The plots show *err* observed at each scalp location for coil orientations of (A) 0°, (B) 45°, (C) 90°, and (D) 135°.

**Figure S5.**
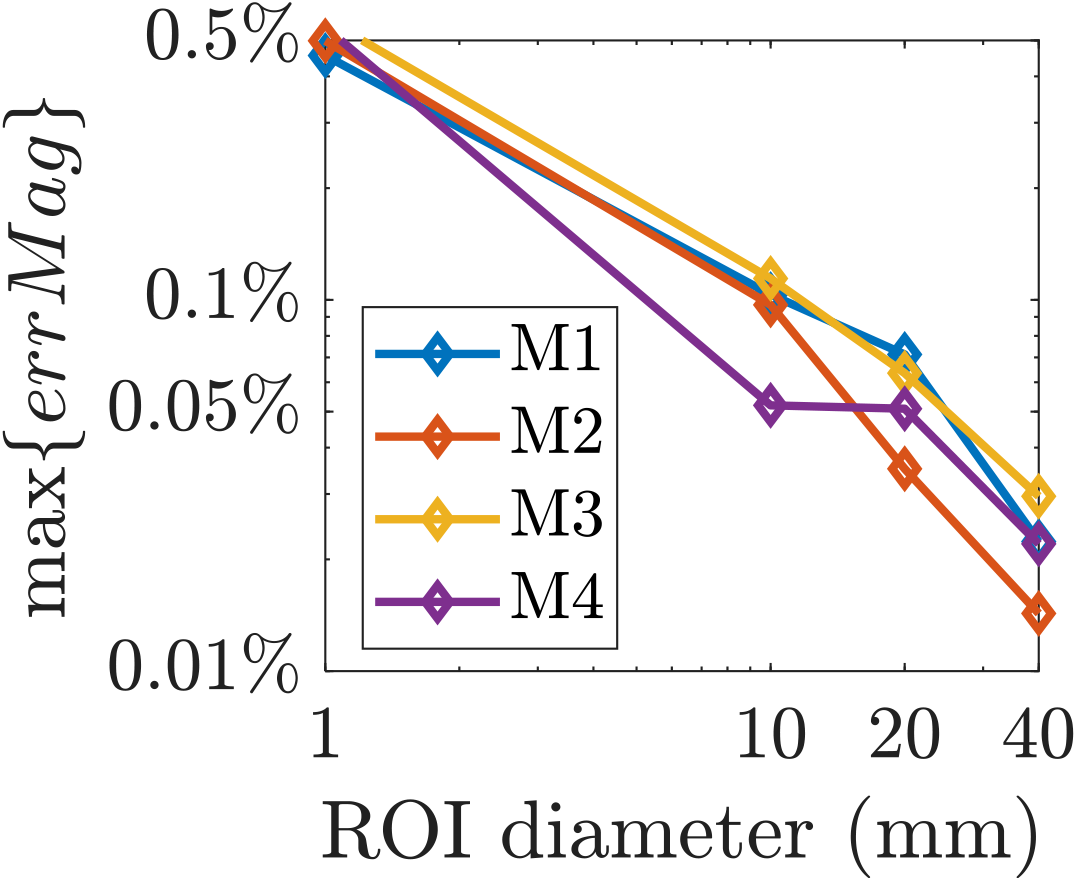
Maximum normalized relative difference for the average E-field magnitude in the SimNIBS simulations compared to our direct FEM results for head models M1–M4.

### Additional coil placement and orientation results

Fig. S6 shows coil placement optimization results analogous to Fig. 8E-T in the main text, but comparing ADM with our in-house direct FEM solver instead of SimNIBS. As in the main results, the optimized placements for the two methods match closely, especially for small ROI sizes.

Tables S1 and S2 provide further quantitative comparisons of the ADM, SimNIBS, and conventional ROI center of mass placement. In Table S1, we observe that the optimized E-field positions deliver improved E-field relative to the coil placement above the ROI center of mass. Futhermore, the optimized coil orientation and the orientation normal to the sulcal wall can differ by as much as 62°. In Tables S2 we additionally compare 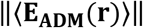 and 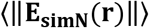 for ROIs of varying sizes. For ROIs with diameter of 20 cm or less the relative difference of the maximum is less than 0.7%. This is below the numerical errors of the FEM solver [1].

**Figure S6.**
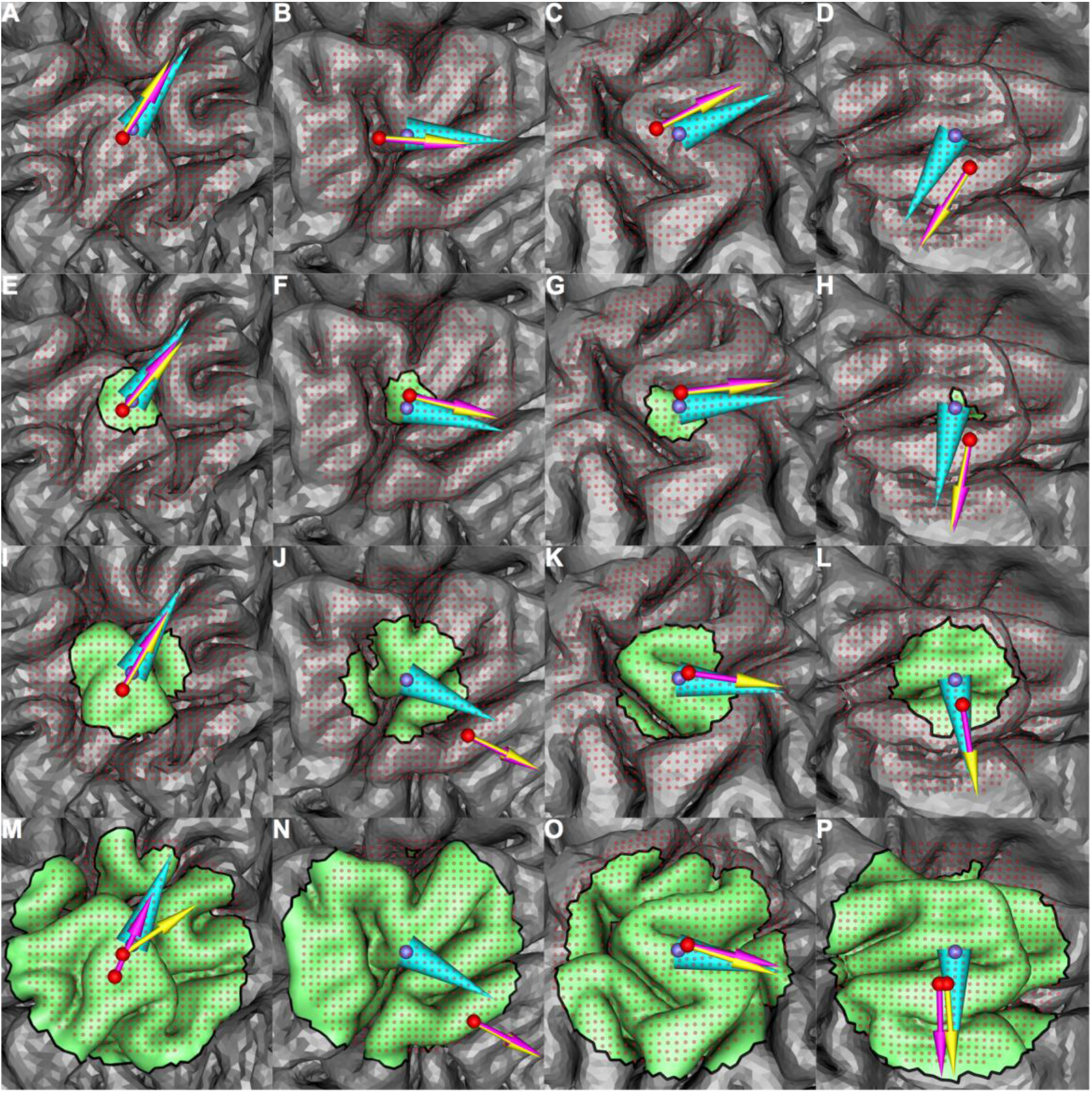
Coil placement results for models M1–M4 (left to right columns) for ROIs of increasing size (top to bottom rows). The in-house direct FEM optimized and ADM optimized position and orientation are represented by the purple and yellow arrow, respectively, 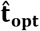 and the ROI center of mass are represented by cyan cone and purple sphere, respectively. The rows show, top to bottom, the results for ROI diameter of 1 (1 tetrahedron), 10, 20, and 40 mm. In most cases both optimization methods result in the same or similar coil position and orientation.

**Table S1:**
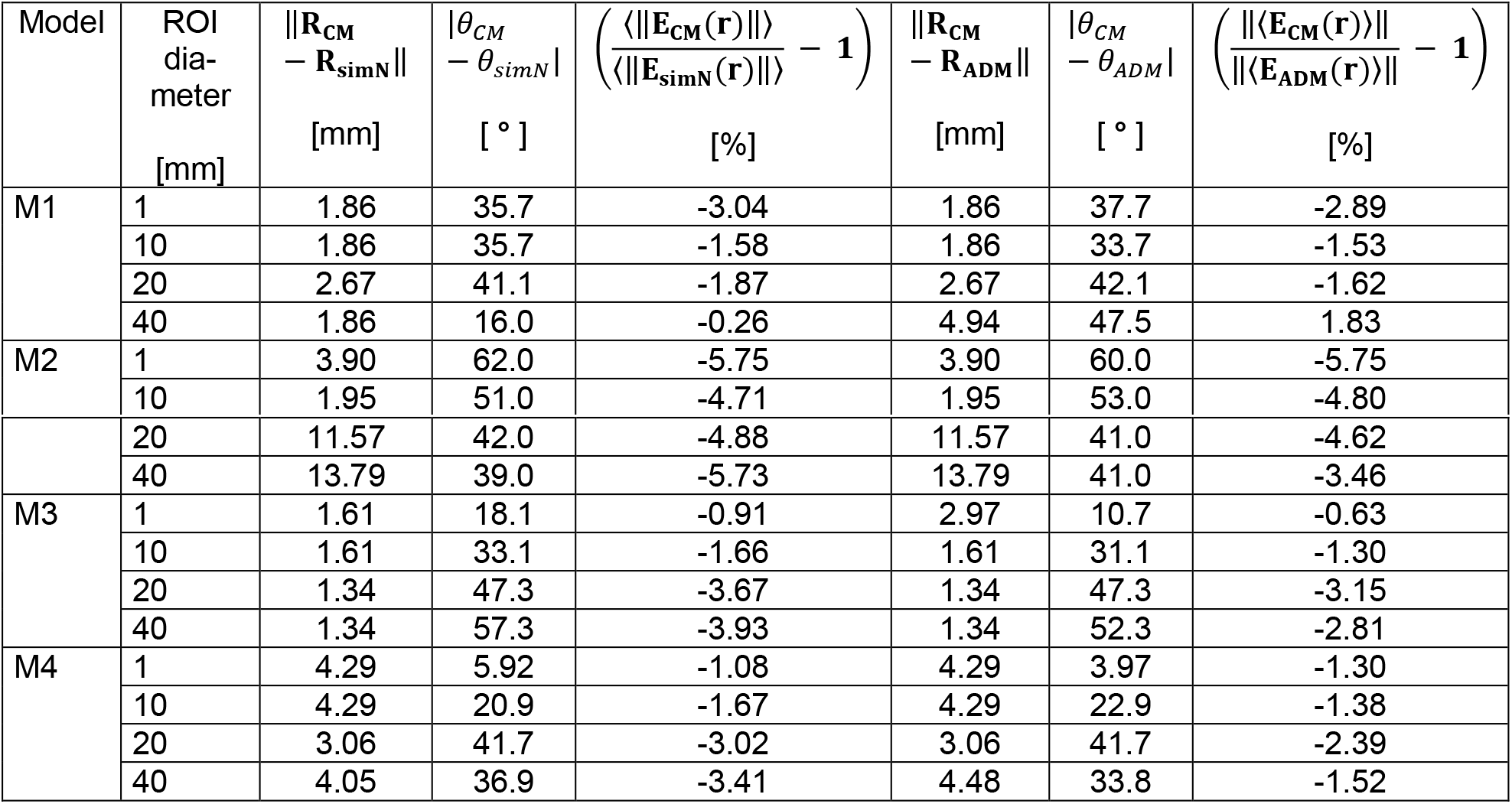
Comparisons between placing the coil above the ROI center of mass (CM) versus where it maximizes the ROI average E-field magnitude with the SimNIBS or ADM methods. Note that the average E-field magnitude is computed differently for the two optimization methods: 〈||·||〉 for SimNIBS comparisons and ||〈·〉|| for ADM comparisons.

**Table S2:** Comparisons between using SimNIBS to maximize the E-field magnitude (i.e. maximize 〈||·||〉) and using ADM to approximately maximize the E-field magnitude (i.e. maximize ||〈·〉||). Unlike in the main text and Table S1, we compare the methods using their respective ROI E-field metrics to emphasize that the E-field magnitude approximation ||〈·〉|| gives a good estimate of the actual average E-field magnitude.

| Model | ROI diameter<br>[mm] | $\ \mathbf{R}_{ADM} - \mathbf{R}_{simN}\ $<br>[mm] | $\vert \theta_{ADM} - \theta_{simN}\vert $<br>[ ° ] | $\left( \frac{\ \langle \mathbf{E}_{ADM}(\mathbf{r}) \rangle\ }{\langle \ \mathbf{E}_{simN}(\mathbf{r})\ \rangle} - 1 \right)$<br>[%] |
| --- | --- | --- | --- | --- |
| M1 | 1 | 0.00 | 2 | -0.15 |
|  | 10 | 0.00 | 2 | -0.049 |
|  | 20 | 0.00 | 1 | -0.25 |
|  | 40 | 3.20 | 32 | -2.1 |
| M2 | 1 | 0.00 | 2 | -0.0035 |
|  | 10 | 0.00 | 2 | 0.095 |
|  | 20 | 0.00 | 1 | -0.28 |
|  | 40 | 0.00 | 2 | -2.4 |
| M3 | 1 | 3.33 | 7.6 | -0.28 |
|  | 10 | 0.00 | 2 | -0.36 |
|  | 20 | 0.00 | 0 | -0.54 |
|  | 40 | 0.00 | 5 | -1.2 |
| M4 | 1 | 0.00 | 4 | 0.23 |
|  | 10 | 0.00 | 2 | -0.3 |
|  | 20 | 0.00 | 0 | -0.65 |
|  | 40 | 1.00 | 3.25 | -1.9 |

## Notes

https://github.com/luisgo/Auxiliary_dipole_method

### Competing Interest Statement

The authors have declared no competing interest.

